# Global Epigenetic Analysis Reveals H3K27 Methylation as a Mediator of Double Strand Break Repair

**DOI:** 10.1101/2021.09.20.461136

**Authors:** Julian Lutze, Donald Wolfgeher, Stephen J. Kron

## Abstract

The majority of cancer patients is treated with ionizing radiation (IR), a relatively safe and effective treatment considered to target tumors by inducing DNA double strand breaks (DSBs). Despite clinical interest in increasing the efficacy of IR by preventing successful DSB repair, few effective radio-adjuvant therapies exist. Extensive literature suggests that chromatin modifiers play a role in the DSB repair and thus may represent a novel class of radiosensitizers. Indeed, chromatin has both local and global impacts on DSB formation, recognition of breaks, checkpoint signaling, recruitment of repair factors, and timely DSB resolution, suggesting that epigenetic deregulation in cancer may impact the efficacy of radiotherapy. Here, using tandem mass spectrometry proteomics to analyze global patterns of histone modification in MCF7 breast cancer cells following IR exposure, we find significant and long-lasting changes to the epigenome. Our results confirm that H3K27 trimethylation (H3K27me3), best known for mediating gene repression and regulating cell fate, increases after IR. H3K27me3 changes rapidly, accumulating at sites of DNA damage. Inhibitors of the Polycomb related complex subunit and H3K27 methyltransferase EZH2 confirm that H3K27me3 is necessary for DNA damage recognition and cell survival after IR. These studies provide an argument for evaluating EZH2 as a radiosensitization target and H3K27me3 as a marker for radiation response in cancer. Proteomic data are available via ProteomeXchange with identifier PXD019388.

## Introduction

Ionizing radiation (IR) remains one of the most widely utilized treatments for cancer, irrespective of organ site or disease stage^1,2^. Modern clinical irradiators can deliver ablative IR doses precisely to the tumor volume while sparing adjacent normal tissue. Even so, radiotherapy is typically ineffective on its own^3,4^ and overcoming resistance often depends on combinations with cytotoxic chemotherapy, which incur greater toxicity^5–7^. An attractive alternative is to identify agents that can enhance local effects of IR on tumor cells but have minimal impacts on unirradiated, normal tissue^8,9^. Although promising candidate radiosensitizers have been identified, failures in translation to the clinic highlight gaps in understanding of radiation response and mechanisms underpinning radiosensitization targets^10–12^. Here, we add to a growing body of literature which establishes chromatin and its constituent histones as key mediators of DNA damage response (DDR) after IR^13^. We further show that modulation of readers and writers of histone post-translational modifications is sufficient to disrupt the cellular response to IR, thereby uncovering additional radiosensitization targets.

Most radiation damage is to DNA bases or single strands, but when IR-induced free radicals react with both strands of chromosomal DNA in proximity, the result can be a double strand break (DSB). DSBs are acutely lethal to cells and are considered the key mediator of radiation’s therapeutic effects; failure of DSB repair can lead to chromosomal instability or aneuploidy ^14^. However, as direct reporters of DSB intracellular location are not yet available, the extent of DSBs is commonly assayed via recruitment of proxy proteins or histone modifications. Prior to any repair, DNA damage must be detected, and break loci marked to direct recruitment of signaling and repair factors. The Phosphatidylinositol 3-kinase-related kinases (PIKKs) ataxia telangiectasia mutated (ATM), Ataxia telangiectasia and Rad3 related (ATR) and DNA-dependent protein kinase (DNA-PKcs) are early responders recruited to DSB loci to mark nearby histone H2AX by phosphorylating Ser139 to form γH2AX^15,16^. γH2AX forms punctate intracellular foci termed IR induced foci (IRIF) which are thought to demarcate repair loci^17,18^. In parallel, poly-ADP ribose polymerase 1 (PARP1) and other PARPs bind proximal to DNA breaks and subsequently PAR-ylate histones and other local substrates, perhaps to affect chromatin decondensation^19–21^. Following break recognition, DSB rejoining is classically described as a choice between two pathways, non-homologous end joining (NHEJ) and homologous recombination (HR)^22,23^. Together these pathways collaborate to restore DNA integrity and limit chromosomal instability^24–27^.

It is widely recognized that chromatin and histone post-translational modifications (PTMs) beyond H2AX phosphorylation impinge upon recognition of DSBs and direct deposition of γH2AX prior to break repair. A range of epigenetic reader and writer enzymes, previously established as transcriptional regulators, have also been implicated in DSB sensing, signaling and repair^28–35^. However, many of these studies are subject to the caveat that DSB formation after IR is dramatically affected by chromatin state. Chromosomal DNA packaged into heterochromatin is intrinsically radiation-resistant compared to actively transcribed DNA^36–39^. Experiments examining epigenetic regulators in DNA damage response typically lack the temporal resolution to distinguish effects on DSB formation from detection or repair. Further, without a direct reporter of DSBs, deconvoluting effects of histone modifiers on IRIF formation or resolution is challenging.

An illustrative example is enhancer of zeste homologue 2 (EZH2), a catalytic subunit of the polycomb repressive complex 2 (PRC2). EZH2 modifies histone H3 to form H3K27me3, a repressive histone mark associated with heterochromatin. PRC2 plays a central role in development, gene silencing, and cell fate decisions via selective deposition of H3K27me3 to assemble heterochromatin^40–43^. As heterochromatin is intrinsically radioresistant, inhibiting EZH2 for one or more cell cycles may phenocopy radiosensitizers by increasing the yield of DSBs without affecting recruitment of DDR factors or recognition of damage. However, EZH2 and other PRC2 subunits also localize to DSBs^44,45^ where EZH2 may deposit H3K27me3 on DSB-proximal nucleosomes^46^. Blocking EZH2 activity immediately prior to irradiation can delay DSB repair, apparently by slowing NHEJ, though inhibiting EZH2 also leads to increased γH2AX levels 24 h after IR insult when NHEJ is no longer thought to participate in repair^47–49^. EZH2 also methylates non-histone substrates and interacts with other DNA damage response factors, adding further complexity ^50–52^. Along with concerns about conflating local and global effects, the notion that a histone mark which mediates heterochromatin contributes to DSB repair is paradoxical as successful repair requires recruitment of several factors and access to DNA.

Development of radiosensitizers has heretofore focused on processes downstream of break recognition such as cell cycle disruption and cell fate. However, a focus on global signaling belies chromatin localized steps critical to DSB repair. The complexity of DSB repair across a varied epigenome coupled with imprecise measurements of DSB repair render radiosensitizer identification challenging. It is thought that small molecules targeting DSB detection by PIKKs or PARPs sensitize cell lines and tumor models to radiation by preventing DSB repair, but clinical translation has lagged ^53,54^. Further, inhibitors of epigenetic readers and writers appear to be attractive radiosensitization targets, but a fuller understanding of their mechanism of action is needed before they can be used clinically.

Toward identifying epigenetic marks that are modulated by DSBs, we used targeted proteomics to evaluate the dynamics of several dozen histone modifications in total chromatin following irradiation. Based on patterns of regulation, this broad survey pointed back to EZH2 as a critical regulator. Toward validating these findings, we confirmed a role for PRC2 in DSB recognition and showed that deregulation of H3K27 modification impacts cellular responses to IR.

## Experimental Procedures

### Cell Culture

MCF7 cells were grown in DMEM medium supplemented with 10% fetal bovine serum (Atlanta) and 4mM L-Glutamine, in a humidified atmosphere of 5% CO_2_ maintained at 37°C. All cells were originally obtained from the American Type Culture Collection (ATCC). The cells were tested for mycoplasma contamination and authenticated by short tandem repeat profile (IDEXX, BioResearch) prior to performing experiments. All experiments were performed within 3 to 10 passages after thawing cells.

### Antibodies

Antibodies used for immunofluorescence in this study are as follows. γH2AX (mouse mAb, clone JBW301, Millipore Sigma) histone H3K27me3 (rabbit, mAb, clone C36B11, CST) histone H3 (mouse, mAb, clone 6.6.2, Millipore Sigma), RNA Polymerase 2 (rabbit polyclonal) R-Loop (Abcam, Rabbit mAb clone S9.6). Secondary antibodies are sheep anti-mouse, Alexa Fluor 488, goat anti-rabbit, Alexa Fluor 647 and Alexa Fluor 595, all sourced from Jackson Immunoresearch.

### Inhibitors and drug treatment

Small molecule probes used in this study were GSK126, an EZH2 inhibitor, and GSKJ4 HCL, a JMJD2/3 inhibitor (Selleck Chem) and veliparib, a PARP inhibitor (obtained from Abbvie). Inhibitor stocks were diluted to 10 mM in DMSO and added to cells for the indicated length of time. Unless otherwise noted, final concentrations used were as follows: GSK126, 20 μM; GSKJ4, 10 μM; Veliparib 10 μM. DMSO was used at 1:1000 dilution for vehicle treatments.

### DNA damage treatment

DNA damage was induced by exposure to a ^60^Co γ-ray source. Cells were placed in an irradiator (MDS Nordion) and exposed to the indicated dose. Dosage rates varied between 10.5 and 9.1 cGy/s depending on the date of the experiment. Cells were allowed to recover in incubator for the indicated time. Non-irradiated (NIR) samples were mock irradiated.

### Multiple Reaction Monitoring (MRM) analysis of histone PTMs

Initial histone PTM analysis was performed by the Northwestern University Proteomics Core. We used the Epiproteomic Histone Modification Panel B assay. The method, in brief, is as follows. Histones were extracted directly from flash frozen cell pellets with the addition of 5 volumes of 0.2 M H_2_SO_4_ for 1 h at room temperature (RT). Cellular debris was removed by centrifugation at 4,000 x g for 5 min and histones were precipitated from the supernatant with trichloroacetic acid (TCA) at a final concentration of 20% (v/v) for 1 h on ice. Precipitated histones were pelleted at 10,000 x g for 5 minutes, washed once with 0.1% HCl in acetone then twice with 100% acetone with centrifugation at 15,000 x g for 5 minutes. After the final acetone wash, histones were dried briefly and stored at −20 °C until derivatization. Histones were propionylated and digested according to Garcia et al. ^55^, with the modification of a single round of propionylation for 1 h prior to and following digestion. Targeted MRM LC-MS/MS was performed on a TSQ Quantiva (Thermo Scientific) triple quadrupole mass spectrometer. This Histone PTM MRM panel B, assaying 95 modification states and their transitions, was developed and setup at the Northwestern Proteomics core and raw data analyzed in Skyline 2 according to published methods^56^.

### Histone PTM data analysis

Data obtained from the MRM analysis were provided as a rectangular matrix with each row representing a PTM and each column containing either the raw peak area (peptide intensity value) or the residue-normalized percentage of a given PTM in a given sample. tSNE analysis was performed in R with the RtNSE package on the residue-normalized data. Default settings were used, though the perplexity was set to 1 because of the low number of datapoints. For clustering of PTMs, raw peak area data were used. The package dtwclust was utilized to perform the DTW distance calculations to obtain more accurate relationships between time-series data. The number of clusters was set at 5 after manual inspection of the elbow plot generated by dtwclust and clustering was carried out via the partitioning around medoids (PAM) algorithm. A heatmap was created using the heatmap2 package in R with a Euclidean distance metric and a Ward D2 clustering algorithm. All plots were generated using ggplot2 implemented in base R or the tidyverse packages. All code used to generate the figures is available upon request.

### Histone sample preparation for LC-MS/MS

Briefly, 5 × 10^6^ cells were harvested, and nuclei were isolated using NEB buffer (10 mM HEPES pH 7.9, 1 mM KCl, 1.5 mM MgCl_2_, 1mM DTT). Histones were extracted from nuclei by treatment with 0.4 N H_2_SO_4_ (Sigma 258105-500mL) for 30 minutes at room temperature and then precipitated from the supernatant by dropwise addition of ice-cold trichloroacetic acid (Sigma T069-100mL). Precipitated protein was spun down and washed twice with very-cold acetone (Fisher A18-500). The pellet was then air dried and resuspended in ddH_2_O. For each sample set, 20 μg of protein (determined via Bradford assay), was loaded and run into a MOPS (Thermo, NuPage NP0001) buffered a 1D 12% gel plug (Thermo NP0341BOX) for 6 min at 200 V.

Gel sections were subjected to propionyl derivatization (at the protein level), Trypsin digestion, propionyl derivatization (at the peptide level), followed by C18 cleanup. For propionyl derivatization, propionic anhydride (Sigma 240311-50g) was mixed 1:3 with isopropanol (ACROS 42383-0040) pH 8.0 and reacted 37 °C for 15 minutes. Following protein derivatization treatment, gel sections were washed in dH_2_O and de-stained using 100 mM NH_4_HCO_3_ (Sigma 285099) pH 7.5 in 50% acetonitrile (Fisher A998SK-4). A reduction step was performed by addition of 100 μl 50 mM NH_4_HCO_3_ pH 7.5 and 10 μl of 200 mM tris(2-carboxyethyl) phosphine HCl (Sigma C4706-2G) at 37 °C for 30 min. The proteins were alkylated by addition of 100 μl of 50 mM iodoacetamide (Sigma RPN6320V) prepared fresh in 50 mM NH_4_HCO_3_ pH 7.5 buffer and allowed to react in the dark at 20 °C for 30 minutes. Gel sections were washed in water, then acetonitrile.

Trypsin digestion was carried out overnight at 37 °C with 1:50-1:100 enzyme– protein ratio of sequencing grade-modified trypsin (Promega V5111) in 50 mM NH_4_HCO_3_ pH 7.5, and 20 mM CaCl_2_ (Sigma C-1016). Peptides were extracted with 5% formic acid (Sigma F0507-1L), then 5% formic acid with 75% ACN, combined and vacuum dried. Post-digestion, peptides were derivatized with propionic anhydride:IPA 1:3 at 37 °C for 15 min and repeated for a total of two times. Peptides were then cleaned up with C18 spin columns (Thermo 89870). and sent to the Mayo Clinic Medical Genome Facility Proteomics Core for HPLC and LC-MS/MS data acquisition via Q-Exactive Orbitrap (Thermo).

### LC-MS/MS and PTM analysis via EpiProfile and MaxQuant

Peptide samples were re-suspended in Burdick & Jackson HPLC-grade water containing 0.2% formic acid (Fluka 94318-50ML), 0.1% TFA (Pierce 28903), and 0.002% Zwittergent 3–16 (Calbiochem 14933-09-6), a sulfobetaine detergent that contributes the following distinct peaks at the end of chromatograms: MH^+^ at 392, and in-source dimer [2 M + H^+^] at 783, and some minor impurities of Zwittergent 3-12 seen as MH^+^ at 336. The peptide samples were loaded to a 0.25uL OptiPak trap (Optimize Technologies, Oregon City, OR) custom-packed with 5um Magic C18-AQ (Michrom BioResources, Inc., Auburn, CA). washed, then switched in-line with a nanoLC column ~34cm × 100um i.d. PicoFrit column (New Objective, Woburn, MA) self-packed with Agilent Poroshell 120S ES-C18, 2.7 um stationary phase. Column flow was 400 nl/min. Mobile phase A was water/acetonitrile/formic acid (98/2/0.2) and mobile phase B was acetonitrile/isopropanol/water/formic acid (80/10/10/0.2). Using a flow rate of 350 nl/min, a 90 min, 2-step LC gradient was run from 5% B to 50% B in 60 min, followed by 50%–95% B over the next 10 min, hold 10 min at 95% B, back to starting conditions and re-equilibrated.

Electrospray tandem mass spectrometry (LC-MS/MS) was performed at the Mayo Clinic Proteomics Core on a Thermo Q-Exactive Orbitrap mass spectrometer, using a 70,000 RP survey scan in profile mode, m/z 340–2000 Da, with lockmasses, followed by 20 MSMS HCD fragmentation scans at 17,500 resolution on doubly and triply charged precursors. Single charged ions were excluded, and ions selected for MS/MS were placed on an exclusion list for 60 seconds. An inclusion list (generated with in-house software) consisting of expected histone PTMs was used during the LC-MS/MS runs.

For EpiProfile analysis, sample *.raw files were extracted and peak picking performed using with pXtract version 2.0 to obtain their MS1 and MS2 files^57^. These along with their *.raw files were analyzed in Matlab with the Epiprofile 2.0 script^58,59^. In addition to the Epiprofile modifications detected, we wanted to probe for any additional common and unique modifications, thus sample *.raw files were also searched in Maxquant version 1.5.2.8 (peaks picked in MaxQuant) against a histone protein fasta database downloaded 10/15/2019 from Uniprot. The PTM search was done in multiple searches at 20ppm with 1% FDR filtering using a fixed modification of Carbamiodomethyl (C), common variable modifications of Deamidation (NQ), Formyl (n-term) Oxidation (M), combined with the following additional PTMS {**Ac** Acetylation (K,S,T), **Ar** ADP ribosylation (R,E,S), **Bu** Butyrylation (K), **Cit** Citruillination (R), **Cr** Crontonylation (K), **Fo** Formylation (K), **Hib** 2-Hydroxyl-isobutyrylation (K), **Ma** Malonylation (K), **Me** Methylation (K,R), **Me2** Di-Methylation (K,R), **Me3** Tri-Methylation (K,R), **Og** O-glycacylation (S,T), **Oh** Hydroxylation (Y), **Ph** Phosphorylation (S,T,Y), **Pr** Propionylation (K), **Su** Succinylation (K), and **Ub** Ubituitylation aka GlyGly (K)}. Downstream PTM analysis was performed in Perseus version 1.6.7.0^60^ and formatted in Perseus, Excel (Microsoft) or R. All code is available upon request. All mass spectrometry proteomics data have been deposited to the ProteomeXchange Consortium via the PRIDE partner repository with the dataset identifier PXD019388^61,62^.

### Immunofluorescence imaging and foci analysis

For all imaging, 2.5 × 10^4^ MCF7 cells were seeded on round #1.5 cover glass in 24 well plates and incubated until 50-80% confluency was achieved. Irradiation and/or treatment with indicated inhibitors were performed *in situ*. For slide preparation, cells were fixed with 4% PFA in PBS for 10 minutes at the indicated time point, stained with 0.5 μg/mL DAPI, and mounted using ProLong Gold (Invitrogen). For immunofluorescence staining, cells were fixed as above, then permeabilized with 10% Triton-X 100 for 10 minutes. After blocking with 5% BSA (American Scientific) in PBS for 1 h, the indicated primary antibodies were added and coverslips were incubated overnight at 4°C. All antibodies were used at 1:1000 dilution. Following three 5 minute washes with 5% BSA in PBS supplemented with 0.1% TX-100 and 0.05% NP-40, fluorescent secondary antibodies (Jackson ImmunoResearch) were applied for 1 h at RT. Foci images were captured on an Olympus IX81 wide-field microscope with either a 40 X or 100 X oil-immersion objective and pseudo colored using ImageJ. Two or more replicates were performed for each experiment and greater than 50 cells were imaged per replicate.

Foci counting was performed with a custom ImageJ macro. Briefly, nuclei were thresholded and segmented and foci were counted within each nucleus via a thresholding and FindMaxima routine. Foci intensity analysis was performed by segmenting the foci as above and then measuring the MFI within each focus. Foci size was determined by auto-local thresholding of the γH2AX channel followed by segmentation and measurement of segmented foci regions. All other image analysis was carried out in ImageJ via custom macros. All macros available upon request.

### Incucyte analysis

For analysis of cellular growth kinetics, MCF7 cells were seeded at low density (10% confluency) in 12-well plates and then treated as indicated. Plates were incubated in the Incucyte S3 imaging system (Essen Biosciences) for 5 days and images were recorded every 4 h. Confluency was calculated automatically using Incucyte software by manually thresholding a random selection of images and applying these settings to the entire image-set. Data were then normalized to the confluency at time of treatment. Plots were generated in R.

### Comet single cell electrophoresis assay

MCF7 cells were irradiated and/or drug-treated as indicated before collection via trypsin and embedding in low-melting agarose (Trevigen). Comet assay was performed with a Trevigen Comet Kit according to manufacturer’s directions with the following modifications. Cells were electrophoresed at 23 V for 60 min and stained with SYBR Green rather than SYBR Gold. Imaging of comet slides was carried out on a wide-field microscope with a 10 X air objective. Images were analyzed using ImageJ plugin OpenComet^63^.

### SA-βGal assay

Cells were seeded at 3×10^4^ cells per well in six-well plates and treated with inhibitors for 1 h prior to irradiation. Cells were allowed to recover in a humidified incubator for 3 days before fixation and staining. Images were captured on a Zeiss Axiovert 200M microscope with a 20× Plan-NeoFluar objective and Axiocam digital camera controlled by OpenLab software. Two or more replicates were performed, and representative images are shown.

### Colocalization analysis

Colocalization between two channels was determined by in-house code written to implement Li’s ICA method^64^. Briefly, ROIs corresponding to individual nuclei were segmented and cropped and images were saved as intensity matrices. A custom R script was written to transform corresponding matrices into colocalization scores. Pixels were considered to be colocalized if the intensity in a given pixel was above the mean intensity for an image in both channels. We reported the fraction of pixels within a given nuclear ROI which were colocalized. This method is insensitive both to the amount of staining present in an image and also to variations in intensity between cells or regions of an image.

### Ground State Depletion (GSD) superresolution imaging

For superresolution imaging, cells were seeded on coverslips and stained as above but not mounted. Coverslips were washed 5X with PBS to remove non-specifically bound fluorophores, inverted over depression slides containing 50 μl of freshly prepared 300 mM MEA oxygen scavenging medium, sealed with a two-part, quick-curing epoxy, and cured 5 minutes in a 50° C oven. For imaging, we utilized a Leica GSD 3D imaging system equipped with a 160 X/1.43 NA, 0.07 mm WD objective; Suppressed Motion (SuMo) stage; PiFoc precision focusing control system; blue (488 nm), green (532 nm) and red (642 nm) excitation lasers; fluorescein, rhodamine and far-red emission filters and an iXon Ultra EMCCD camera. Slides were then imaged using standard GSD imaging protocols with at least 10,000 frames captured per channel per image. GSD data analysis and processing were carried out with a series of in-house ImageJ macros. Identification of emission events was performed via ImageJ plugin ThunderSTORM^65^. Final images were then pseudo colored and compiled in ImageJ. Superresolution imaging macros are available upon request.

### GSD Fluorescence Resonance Energy Transfer (FRET) imaging

We labeled target proteins or PTMs with primary antibodies as indicated and utilized fluorescent secondary antibodies to introduce either a donor fluorophore (AF 594) or an acceptor fluorophore (AF 647), hereafter referred to as donor (DNR) and acceptor (ACC) respectively. First, both DNR and ACC were imaged at their respective excitation maxima to obtain an image of DNR and ACC location. Following high laser power exposure for 60s, both DNR and ACC were reimaged at their respective excitation maxima. The second ACC image displayed negligible signal indicating efficient bleaching on ACC fluorophores. Before ACC bleach, DNR energy was transferred to the ACC proportionately to the distance between DNR and ACC molecules. Bleached ACC fluorophores can no longer accept DNR energy, and all DNR energy is thus observed when exciting DNR fluorophores at DNR excitation maxima. Any increase in the DNR emission after ACC bleach is thus indicative of FRET and proportionate to the distance between ACC and DNR molecules. To obtain a FRET image, the DNR image before ACC bleach is subtracted from the DNR image after ACC bleach. The resultant image intensity is proportional to FRET between ACC and DNR. GSD-FRET reports both the location and the degree of FRET interactions between two labeled antigens. GSD-FRET imaging was carried out in the sequence described above. Images were pseudo-colored and manipulated in ImageJ.

### Experimental Design and Statistical Rationale

For EMHP analysis, samples were analyzed in technical triplicate (n=3). For EpiProfile validation, biological triplicate samples were also collected (n=3). Number of replicates was selected based on standard proteomic experimental design. For EHMP analysis, peptides were selected in accordance with established protocols. For EpiProfile and MaxQuant analysis, data was generated with LC-MS/MS gradients compatible with EpiProfile, and ions were selected for MS/MS based on the Top20 most abundant peaks or if it appeared on an inclusion list consisting of common histone PTM proteotryptic peptides’ expected ion masses. Inclusion list ions were generated via in-house software selecting for (Ac, Me1-3, Ph or Ub PTMs across major histone isoforms). Peptides were selected using the EpiProfiler software or MaxQuant software according to default parameters. We performed non- controls for all experiments and data was collected in a time-course manner. Samples were not processed in a blinded fashion, though the order in which samples were processed was designed to minimize sample carryover and our protocol includes a blank injection between runs to mitigate column carryover. For EpiProfiler and MaxQuant analysis, raw files were searched at 1% FDR. A MaxQuant peptide cutoff score of 40 was also used for PTM peptide analysis. All statistical analysis was performed as indicated. Test were carried out in R using the ggpubr package. In general, a Wilcox Ranked-Sum Test was used to compare two samples. Kruskal-Wallis tests were used for analyses of more than two groups. For EpiProfile analysis, a Friedman test was performed in Prism (GraphPad). For all plots, significance values are as follows: ns p>0.05; * p<0.05; ** p<0.01; *** p<0.001; **** p<0.0001. Box plots show first and third quartiles of the data as well as the median. In scenarios where multiple testing was considered, p-values were transformed into FDR q-values by the qvalues package in R (Storey method). All plots were generated in R using the ggplot, cowplot and ggpubr packages. Boxplots show the median, 1^st^ and 3^rd^ quartiles with whiskers extending to 1.5*IQR. All software versioning is described in Methods. All R code, for data generation, analysis, and plotting is available upon request.

## Results

### Ionizing radiation induces widespread and long-lasting alterations to histone post-translational modifications

Targeted analyses probing one or a few histone post-translational modifications (PTMs) at a time have revealed a limited set of modifications that regulate the DNA damage response (DDR)^66–68^. However, this work has not examined modifications beyond well-characterized epigenetic marks such as Ac, Me, Ph, and Ub, despite the rapidly growing list of dynamic histone modifications^69,70^. Methods for proteomic analysis of global histone PTMs are now well established and provide broad coverage of most individual and many combinatorial histone modifications^58,59^. Such methods have yet to be fully utilized to track epigenetic changes following genotoxic stress.

Toward surveying a broad range of epigenetic marks, we initially applied a multiple reaction monitoring (MRM)-based targeted quantitative triple-quadrupole mass spectrometry assay, the Epiproteomic Histone Modification Panel (EHMP, Northwestern Proteomics), to analyze multiple histone PTMs over a time course following irradiation of MCF7 breast carcinoma cells using a ^60^Co source. ^60^Co γ rays induce a wide range of DNA lesions, including a high fraction of complex DSBs which are characterized by multiple chemical changes that preclude rapid end-joining ^71^. To sample a time-course spanning DSB formation to the anticipated completion of most repair^72^, MCF7 cells were irradiated with 6 Gy, returned to culture, and samples were collected at 1, 4, 24 and 48 h post IR (PIR). Acid-extracted histones from control and irradiated cells were subjected to the MRM histone PTM survey to measure modifications at a per-residue level. For residues which could be in any one of several modification states such as H3K9 (which may be unmodified, acetyl, mono-, di- or tri-methylated) analysis indicated the fraction of residues in each state. Thereby, 92 histone modifications were evaluated on 30 histone residues for each sample (Supplementary Table 1).

To assess overall changes in PTMs over the time course, the data for three technical replicates for each time point were examined by t-distributed stochastic neighbor embedding (tSNE) (Fig. 1a). Each time point after 6 Gy was distinct from the unirradiated (NIR) control. The 1 h and 4 h PIR samples clustered together and 24 h and 48 h PIR samples formed a separate cluster suggesting IR produces separable short and long-term changes to the epigenome. As a complementary approach, we applied hierarchical clustering (Fig. 1b), yielding relationships between the samples. The samples again fell into distinct groups corresponding to short-term and long-term changes. The shared patterns of PTM dynamics revealed by clustering were analyzed further by dynamic-time warping (DTW) analysis to extract temporally distinct modification trajectories (Fig. 1c). After clustering, five trajectories emerged from the data (Supplementary Table 2). Mapping the cluster centroids of each trajectory indicated a range of histone modification dynamics in response to DNA damage. In particular, Cluster 2 included many of the PTMs that increased sharply by 1 h PIR, including H3K9 methylation, H3K27 methylation, as well as H4 acetylation, known to mediate 53BP1 recruitment^73^.

**Figure 1.**
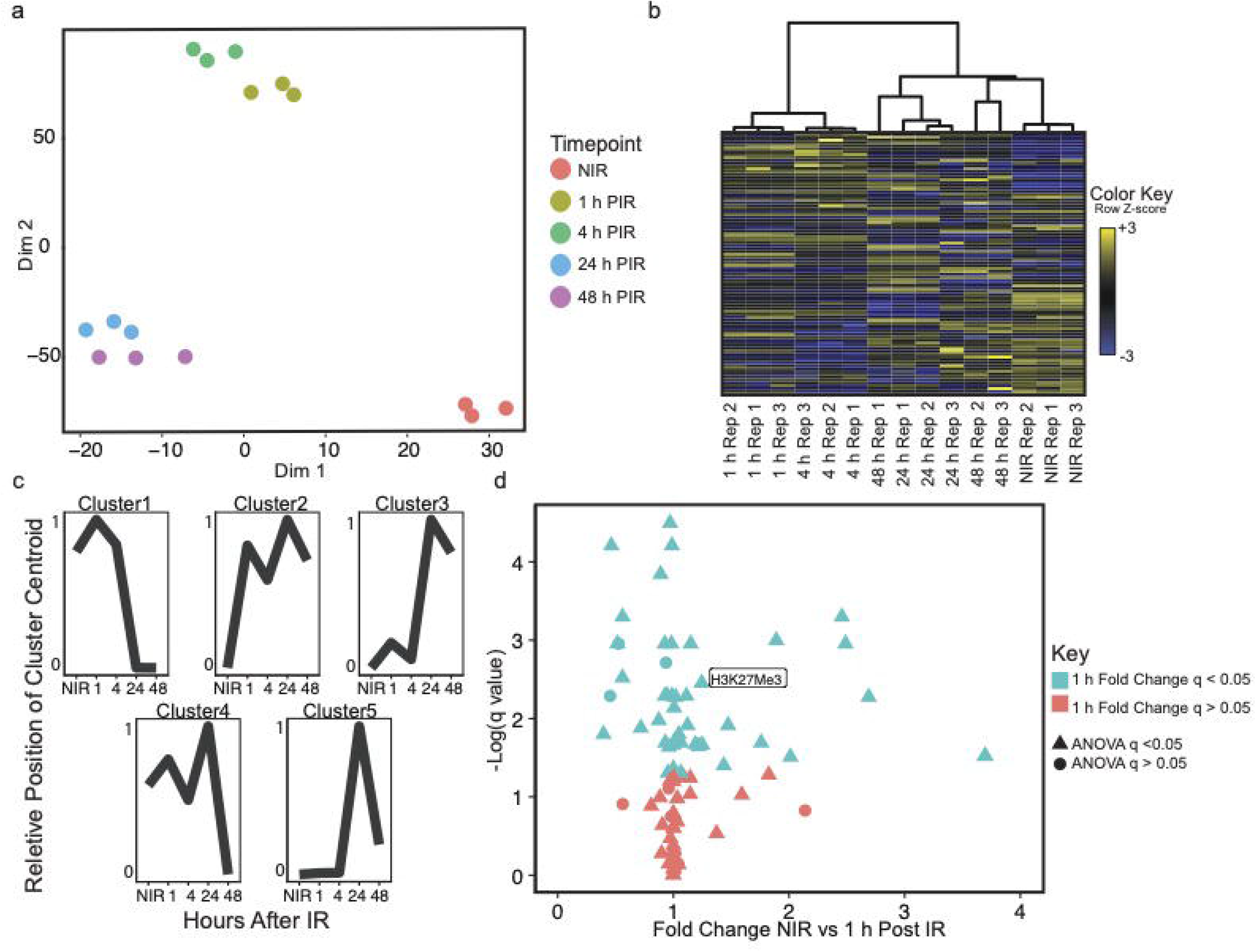
Histone post-translational modifications (PTMs) are dynamically altered by DNA damage induction. a) tSNE of samples from MRM histone PTM time-course analysis. Dots represent samples from technical replicates (*n* = 3), color-coded to denote the timepoint after exposure to mock-irradiated (NIR) or 6 Gy (IR), using a ^60^Co γ-ray source. Data used to compute the tSNE are histone PTM per-residue percentages from the EHMP assay. b) Heatmap of the matrix used to generate the tSNE plot in Fig.1a. Heatmap is clustered by Euclidian distance between samples. Data used are histone PTM per-residue percentages from the EHMP assay. Three replicates are shown. c) Centroid plots of histone PTM clusters. Data used are histone PTM per-residue percentages from the EHMP assay, averaged between three replicates. Average trajectories of all PTMs were clustered according to their Dynamic Time Warping distance and then centroids were fitted and plotted by the PAM algorithm. The number of clusters was set to 5 after manual inspection of the data. The Y-axis denotes the relative average PTM density in each cluster normalized to the NIR timepoint. d) Volcano Plot of all PTMs analyzed. X-axis denotes the average fold change between the NIR and 1 h PIR timepoints. Y-axis shows the negative log of the FDR corrected P-value. Points are color-coded according to their significance at 5% FDR (comparison of NIR to 1 hPIR by Wilcox Ranked-Sum Test) and their shape denotes the significance for a Kruskal-Wallis test across all timepoints, also at 5% FDR. H3K27me3 is labeled for clarity.

Toward identifying specific PTMs involved in the DNA damage response, we examined which modifications were significantly changed over the time course. Of the 92 PTMs evaluated, 78 displayed significant changes at one or more timepoints after correcting for multiple testing (5% FDR; Kruskal-Wallis test) (Supplementary Table 3). Pointing to pathways that may mediate early events such as DNA damage recognition and signaling, 58 PTMs were significantly altered at 1 h PIR compared to non-irradiated cells after correcting for multiple testing (5% FDR; Wilcox Ranked-Sum Test, Fig. 1d) and 51 were both significantly higher at 1 h PIR and dynamic across the time course. This group included several PTMs previously linked to DSB repair including H3K79 methylation, catalyzed by Dot1L, and H4 methylation at K8, K12, K16 or K20, which mediate 53BP1 binding and NHEJ repair (Supplementary Fig 1)^73,74^. Further validating this approach, our analysis identified H3K27 trimethylation (q=0.031, Kruskal-Wallis; FC=1.153, q=0.0078, Wilcox) as significantly increased 1 h PIR. As noted above, H3K27 trimethylation has been linked to DNA damage response and NHEJ^45,48^, but mechanisms remain poorly defined.

### An accurate mass and time approach confirms global epigenetic changes after irradiation

As a complementary approach, we performed an independent time-course analysis of chromatin modifications after irradiation using label-free, conventional LC-MS/MS and data analysis with EpiProfile 2.0^58^, an accurate mass and time (AMT) strategy to quantify over 200 histone marks (Ac, Me1, Me2, Me3, and Ph). Following the EpiProfile protocol, histones from irradiated MCF7 cells were enriched by acidic extraction and then propionyl derivatized before and after trypsin digestion. The resulting peptides were subjected to Orbitrap LC-MS/MS in biological triplicate then examined with EpiProfile, manually validating spectra for PTM sites of interest. This analysis detected 204 PTM combinations reducing down to 45 single PTMs and found dynamic changes in 38 PTMs during the time course (Supplementary Table 4).

Focusing on modifications of histone H3 isoforms, we observed significant changes (Kruskal-Wallis, p<0.05) across several residues including H3K27 and H3K36 (Supplementary Fig 2). We then plotted fold changes for PTMs on H3 residues at each of the four timepoints compared to unirradiated cells. The resulting heatmap revealed kinetically distinct patterns of dynamic modification for specific residues and PTMs including an increase in H3K27 and H3K36 methylation (Fig. 2a). Plotting relative PTM changes as compared to an unirradiated control, grouped by modification type, revealed a significant trend toward increased acetylation and conversion of mono-methylation to di- and tri-methylation, particularly during the first 24 h PIR (Fig. 2b). While effects of histone modifications are residue-specific, a global reduction in acetylation and an increase in methylation may suggest chromatin compaction or gene repression following IR. Next, we separated the H3K27 modification data by H3 isoform. This analysis revealed that H3.3 experiences the bulk of the observed reduction in K27 di-methylation as well as the increase in K27 tri-methylation (Fig. 2c-d). The MRM method was not powered to detect isoform level changes in H3, illustrating the added analytical power of Epiprofiler. H3.3 is enriched in euchromatin; thus, a potential role for increased H3K27me3, a PTM linked to transcriptional silencing, may be to suppress conflict between transcription and repair^75^.

**Figure 2.**
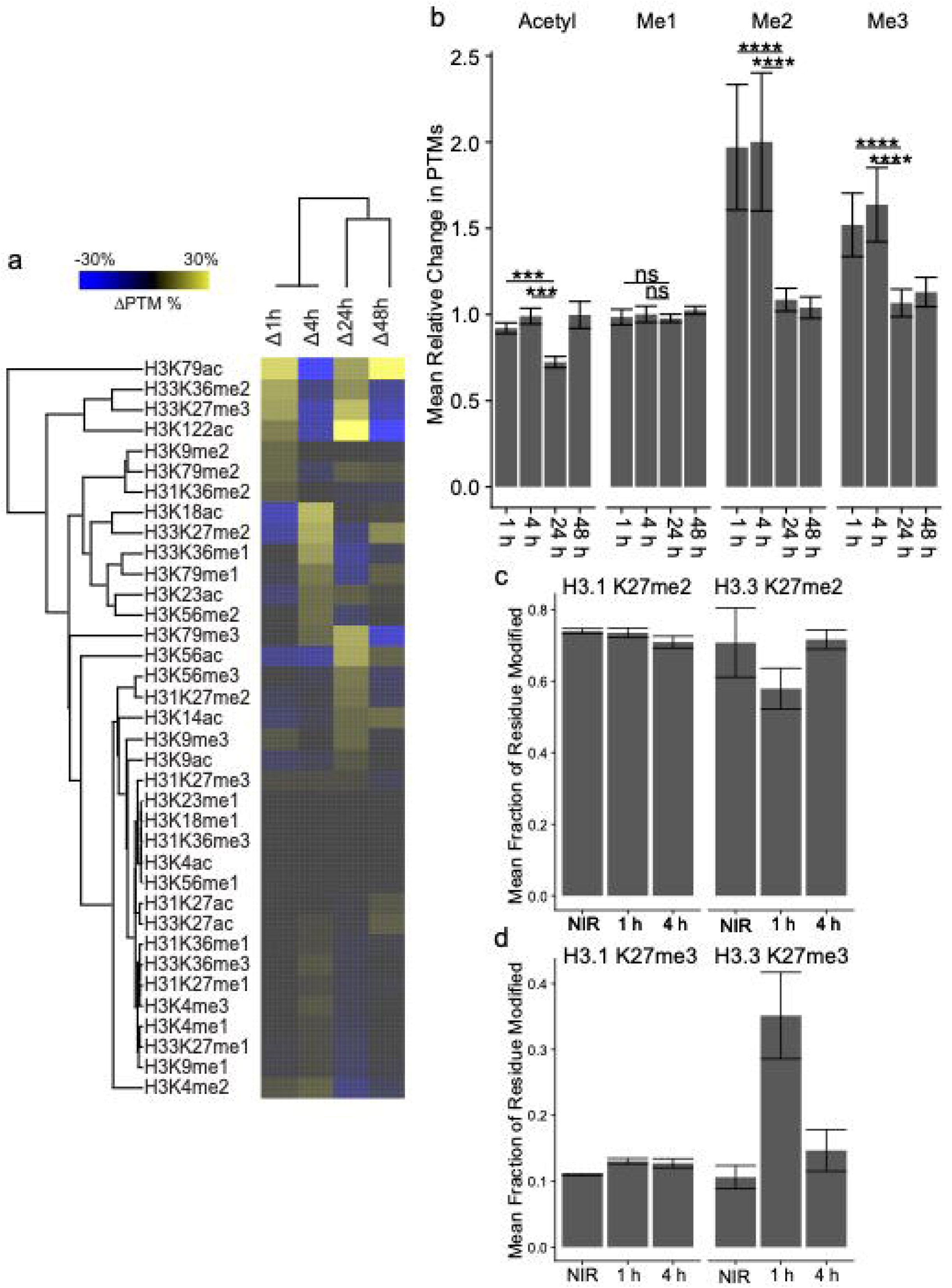
Epiprofile 2.0 quantification of temporal histone marks after DNA damage induction. a) Heatmap of the relative changes between timepoints for PTMs on Histone H3 and Histone H4. Data are the average percent PTM change between NIR samples and the indicated timepoints. Data from biological replicates (*n* = 3). Note the time-point specific regulation of various groups of marks. b) Plot shows average modification changes for acetyl, mono-, di-, and tri-methylation across all residues measured for each of the timepoints relative to NIR. Data from biological replicates (*n* = 3). We observe a decrease in acetylation following IR and an increase in overall methylation specifically me2 and me3 at the 1 h PIR timepoint. Error bars show SEM between average PTM values. Significance was determined by Wilcox Ranked-Sum Test between indicated timepoints. Significance values are as follows: ns p>0.05; * p<0.05; ** p<0.01; *** p<0.001; **** p<0.0001. Total number of PTMs are: Ac-13, me1-13, me2-7, me-3 7. c) Abundance of H3K27 di-methylation separated by H3 isoforms at 1 and 4 h PIR. Plot shows the percent of the residue in each modification state. The magnitude of changes is much larger for H3.3, an isoform associated with euchromatin. Error bars show SEM between average PTM values across biological replicates (*n* = 3). d) Abundance of H3K27 tri-methylation separated by H3 isoforms at 1 and 4 h PIR. Plot shows the percent of the residue in each modification state. The magnitude of changes is much larger for H3.3, an isoform associated with euchromatin. Error bars show SEM between average PTM values across biological replicates (*n* = 3).

Comparing the DDA EpiProfile method to the MRM EHMP panel, and not accounting for unmodified peptides, EpiProfile detects a total of 161 PTMs and EHMP detects 63 PTMs, with 55 PTMs shared between the two methods. Considering only common PTMs, EpiProfile displayed an inter-replicate R^2^ of 0.92 and EHMP an inter-replicate an R^2^ of 0.99 after linear regression analysis. Nonetheless, comparing the two methods yields an R^2^ of only 0.17 (Supplementary Fig 3), likely reflecting distinct biases between the two assays that affect sensitivity toward different PTMs.

### Untargeted analysis reveals additional dynamic histone PTMs during the DNA damage response

Recent work has expanded the universe of histone modifications of importance beyond acetylation, methylation, phosphorylation and ubiquitinoylation with the discovery of novel modifications including new PTMs such as Crontonylation (Cr), Acetylations, O-GlcNAcylation (Og), Propionylation (Pr), Butyrylation (Bu), and ADP ribosylation (Ar)^70^. To extend the analysis beyond the sites identified by EHMP or EpiProfile, the *.RAW data files obtained from QE-Orbitrap LC-MS/MS of the acid extracted and propionylated histone peptides were searched with MaxQuant to detect additional dynamic histone PTMs. We queried the Epiprofile data files for 17 additional PTMs: Ac, Ar, Bu, Cit, Cr, Fo, Hib, Ma, Me1, Me2, Me3, Og, OH, Ph, Pr, Su, Ub. PTMs were split into groups of 3-4 modifications to balance CPU load, search time, and data search space. This analysis detected 3076 total modifications across 5 time-points. (Supplementary Table 5, 6). Summary counts for unique PTMs were as follows, Ac: 516, Ar: 1, Bu: 236, Cit: 28, Cr: 132, Fo: ND, Hib: 64, Ma: 35, Me1: 531, Me2: 95, Me3: 171, Og: 39, OH: ND, Ph: 38, Pr: 956, Su: 28, Ub: 206). The ability to incorporate additional searches on EpiProfile data is an additional advantage over MRM methods such as EHMP. We acknowledge the ability to eventually add these new modifications to the EpiProfile MS based software as a custom search or potential version release. Although novel PTMs warrant further study, they remain challenging to assay experientially as readers and writers are unknown. Thus, we focused on the well characterized modification H3K27me3 which was highlighted by both EHMP and EpiProfile analyses above.

### Inhibition of K3K27 methylation attenuates DSB recognition and repair

H3K27 trimethylation, mediated by the PRC2 catalytic subunit EZH2, is associated with transcriptional repression, heterochromatin formation and maintenance,^76,77^ and is also implicated in DSB detection and NHEJ repair^44,78^. Demethylation by the jumonji-domain demethylases JMJD2 (KDM4A) and JMJD3 (KDM6B) opposes EZH2 activity^79,80^. Toward establishing functional significance of H3K27 methylation in DSB recognition and repair, we acutely exposed MCF7 cells to EZH2 and/or JMJD2/3 inhibitors at tenfold over IC_50_ to ablate enzyme activity prior to irradiation (Fig. 3a). As a control, we inhibited PARP1 with the non-trapping inhibitor veliparib^81^, known to delay DSB repair. Expected effects of each inhibitor on H3K27 methylation were confirmed by immunostaining with an anti-H3K27me3 antibody (Fig. S4a).

**Figure 3.**
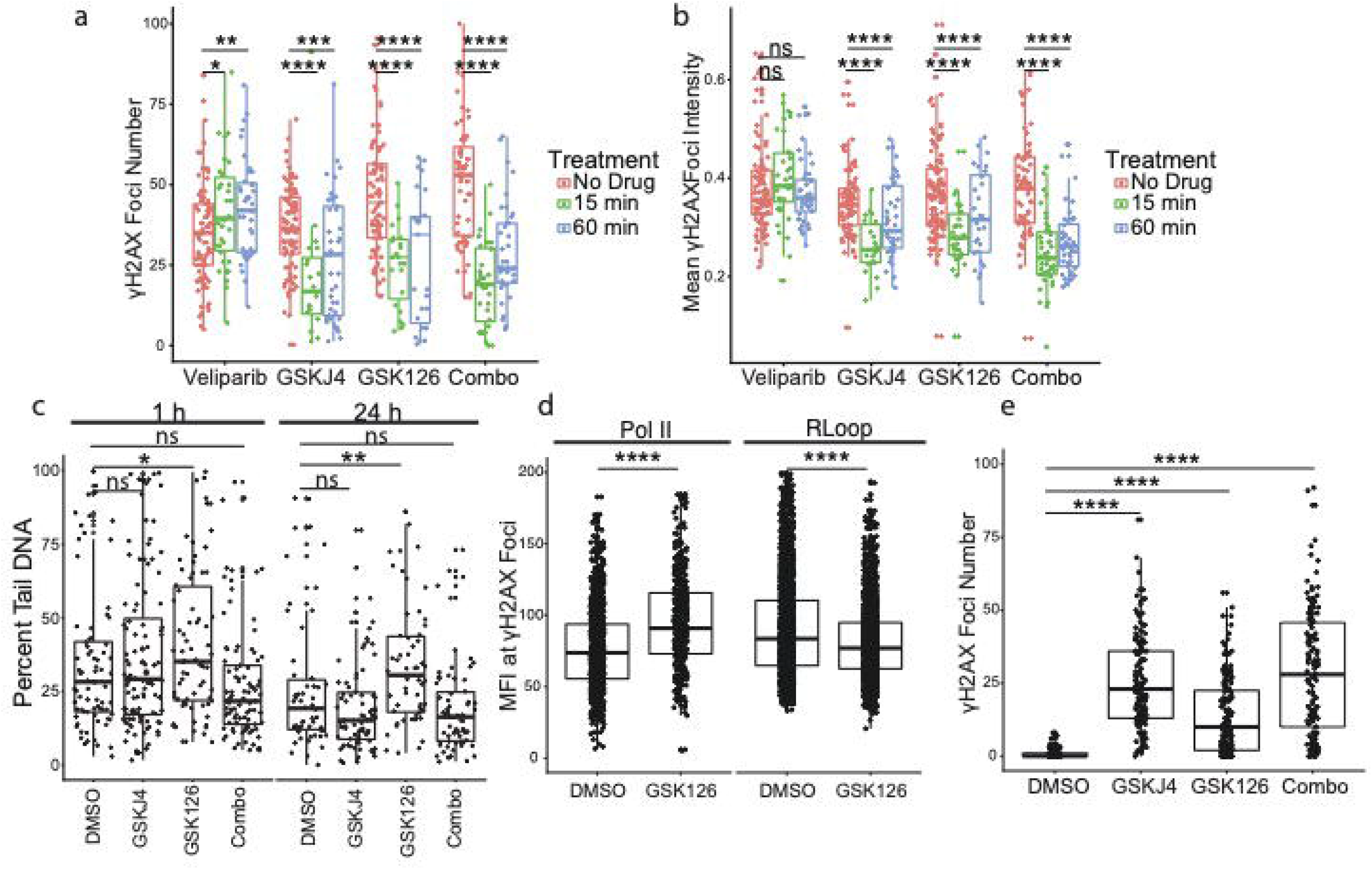
Inhibition of H3K27 methylation attenuates DSB recognition and repair. a) Mean number of γH2AX foci after drug treatment. Foci counting was performed by a custom ImageJ macro. Drugs were added for the indicated length of time prior to dosing with 6Gy of IR. Cells were fixed and stained 1 h PIR. Combo refers to a mixture of both GSK126 and GSKJ4 at their original concentrations. Significance was determined by a Kruskall-Wallace test performed within each treatment group. Significance values are as follows: ns p>0.05; * p<0.05; ** p<0.01; *** p<0.001; **** p<0.0001. Three biological replicates were collected. Total number of points are: 86, 33, 37, 90, 21, 39, 84, 21, 25, 62, 29, 36 per group from left to right. b) Plot as in Fig. 3b but showing the mean γH2AX foci intensity. Foci intensity analysis was performed by a custom ImageJ macro. Three biological replicates were collected. Significance testing as in Fig. 3a. Total number of points are: 86, 33, 37, 90, 21, 39, 84, 21, 25, 62, 29, 36 per group from left to right. c) Comet assay results of cells treated as in Fig. 3b and assayed either 1 or 24 h PIR. Plotted is the Tail DNA percent as reported by the ImageJ plugin OpenComet. Significance was determined by a Wilcox Ranked-Sum Test against DMSO treatment. Three biological replicates were collected. Total number of points are: 78, 103, 67, 94, 55, 80, 52, 68 from left to right. d) Mean fluorescence intensity of the indicated antigens at γH2AX foci. Foci intensity analysis was performed by a custom ImageJ macro. Three biological replicates were collected. Total number of points are as follows: 1210, 416, 2790, 2738 from left to right e) Plot as in Fig. 3a, but performed 24 h PIR. Foci counting was performed by a custom ImageJ macro. Drugs were added for 1 h prior to dosing with 6 Gy of IR and media was exchanged 1 h after IR insult. Three biological replicates were collected. Total number of points are as follows: 144, 142, 151, 149 from left to right

MCF7 cells were treated with the EZH2 inhibitor GSK126 (20 μM) alone or in combination with the JMJD2/3 inhibitor GSKJ4 (10 μM) for the indicated length of time, exposed to 6 Gy of IR, and allowed to recover for 1 h before being fixed and immunostained for γH2AX 1 h PIR. Acute treatment with the inhibitors, alone or in combination, significantly decreased γH2AX foci number after radiation (Fig. 3b). Additionally, we observed a significant reduction in fluorescence intensity of γH2AX foci in cells treated with GSK126 and/or GSKJ4, suggesting that dysregulation of H3K27 methylation limits local H2AX phosphorylation (Fig. 3c). Conversely, treatment with the PARP inhibitor veliparib (10 μM) somewhat increased γH2AX foci number and intensity (Fig. 3b, c). The short interval between drug treatment and irradiation precludes effects dependent on gene repression or chromatin condensation and instead suggests that H3K27 methylation may be necessary for break recognition, as previously described^78^ and that methylation acts upstream of H2AX phosphorylation.

To assess effects on radiation sensitivity, cells were treated with GSK126 alone or in combination with GSKJ4 for 1 h and irradiated with 6 Gy. The media was then replaced to relieve epigenetic inhibition, and cell growth was followed for 5 days by time-lapse imaging in an IncuCyte imaging incubator system. Here, the two drugs yielded different phenotypes: treating cells with the JMJD2/3 inhibitor completely blocked proliferation, while the EZH2 inhibitor attenuated recovery from IR (Fig. S4b). Single cell electrophoresis (comet) assay was performed to assess DSB repair independently from γH2AX foci resolution. At 1 h PIR, we observed an increase in unrepaired DSBs following GSK126 treatment (Fig. 3d). These breaks persisted 24 h PIR, indicating that EZH2 inhibition leads to unrecognized or irreparable damage and loss of genomic stability. Examining the IncuCyte data, we observed both a decrease in proliferation and cell death following GSK126 treatment, likely a consequence of unrepaired DSBs (Fig. S4c). Toward establishing a mechanism by which short-term GSK126 treatment attenuates DDR repair, we examined transcription at DSB loci. In order to prevent additional damage, transcription must be attenuated proximal to broken DNA. R-Loops, a product of stalled transcription, are thought to participate in rapid repair of some DSBs arising in transcriptional units ^82,83^. We examined both RNA Polymerase II (Pol II) and R-Loops at γH2AX foci with or without GSK126 treatment (Fig. 3e). Attenuation of H3K27 methylation resulted in an increase in Pol II and a decrease in R-Loops at γH2AX foci 1 h PIR suggesting transcription is not properly attenuated, perhaps resulting in impeded DSB repair or leading to further damage by transcribing across broken DNA.

We next investigated the dramatic attenuation of proliferation upon GSKJ4 treatment. SA-βgal staining was conducted to assess cell senescence^84^. Persistent DDR signaling can drive cells toward senescence, even in the absence of unrepaired DSBs. Cells were treated as in the IncuCyte experiment and stained for SA-βgal five days post IR exposure (Fig. S4d). These data reveal that GSKJ4 treatment, alone or in combination with EZH2 inhibition, increases cellular senescence. However, GSK126 treatment did not increase senescence. We examined γH2AX foci 24 h PIR in combination with 1 h treatment with GSK126, JMJD2i or a combination. Indeed, exposure to GSKJ4 alone or in combination with GSK126 led to increased persistent γH2AX foci indicative of a failure to wind down damage signaling following end joining. Persistent DSB signaling has been shown to trigger senescence^84,85^. Thus, H3K27me3 may act at multiple stages during the DDR process. Perhaps H3K27me3 deposited at breaks is required for end joining, but must later be removed by JMJD proteins. Failure to do so triggers persistent DDR signaling leading to senescence^86–88^.

### Imaging confirms that H3K27me3 is deposited proximal to DSBs and mediates γH2AX formation

Toward confirming a local effect of H3K27 methylation at DSBs, we examined colocalization of H3K27me3 and γH2AX after irradiation. Conventional immunofluorescence analysis 1 h PIR revealed punctate domains of increased H3K27me3 immunoreactivity along with significant overlap between H3K27me3 and γH2AX (Fig. 4a). Colocalization analysis revealed diminished colocalization of H3K27me3 and γH2AX after treatment with GSKJ4 or GSK126, as compared to vehicle treatment, or addition of the PARP inhibitor veliparib (Fig 4b).

**Figure 4.**
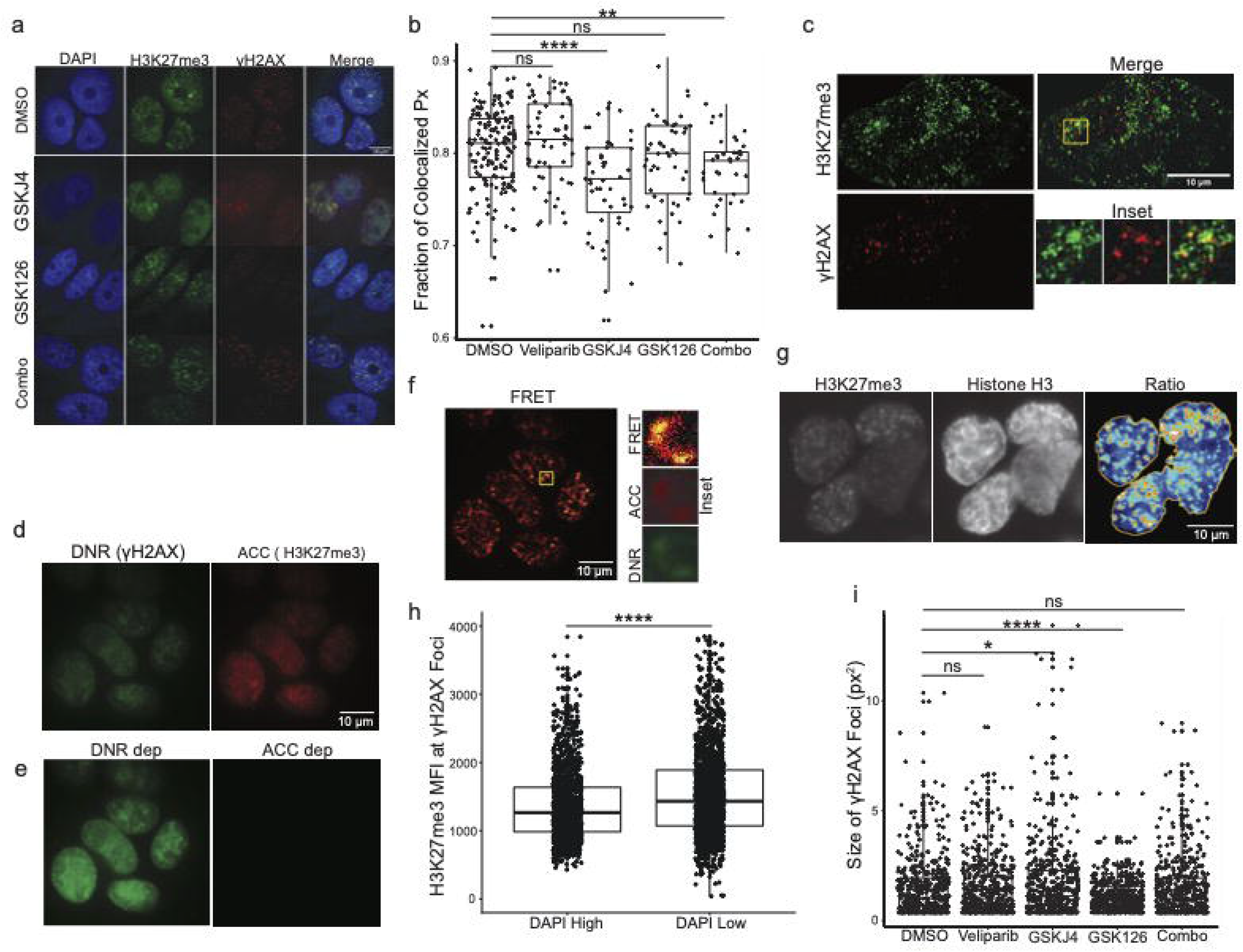
H3K27me3 is a local determinant of DSB recognition. a) Immunofluorescence images of irradiated MCF7 cells. Cells were treated with the indicated drugs for 60 minutes prior to dosing with 6 Gy. Cells were fixed and stained 1 h PIR. Images were acquired using a 40 X oil objective on a spinning-disk confocal microscope. A representative image is shown from 3 replicates. b) Quantification of colocalization between γH2AX and H3K27me3 staining in the slides shown in Fig. 4a. The fraction of colocalized pixels was calculated per nucleus using Li’s ICA method. Significance was determined by a Wilcox Ranked-Sum Test against DMSO treatment. Significance values are as follows: ns p>0.05; * p<0.05; ** p<0.01; *** p<0.001; **** p<0.0001. Three biological replicates were collected. Total number of datapoints are as follows: 165, 56, 45, 41, 31 per group from left to right. c) Superresolution imaging of irradiated MCF7 cells. DMSO treated cells were fixed 1 h PIR and imaged on a Leica GSD imaging system. Inset shows colocalized puncta of H3K27me3 and γH2AX. A representative image is shown from 3 replicates. d) GSD-FRET analysis of colocalization between γH2AX and H3K27me3. DMSO treated cells were fixed 1 h PIR and imaged on a Leica GSD imaging system using a 160x objective. Both Donor and Acceptor channels were imaged at their respective excitation maxima. A representative image is shown from 3 replicates. e) Cells as in Fig. 4d but following depletion of the Acceptor fluorescent dye using intense laser power for 2 minutes. Imaging conditions were equivalent to Fig. 4d. f) Pseudo colored image showing the relative increase in signal in the Donor channel following Acceptor photobleach (Fig. 4e, left minus Fig. 4d, left). Inset shows region with both γH2AX and H3K27me3 signal from panel Fig. 4d alongside the same region from Fig. 4f. g) Ratio-based imaging of irradiated MCF7 cells. Cells were fixed 1 h PIR and imaged using a 40 X oil objective on a spinning-disk confocal microscope. Ratios between channels were calculated in ImageJ by dividing image intensities and then the resulting image was thresholded. Rightmost panel was pseudo-colored to highlight differences in H3K27me3:H3 ratio. A representative image is shown from 3 replicates. h) Mean fluorescence intensity of H3K27me3 at γH2AX foci after GSK126 treatment. Drugs were added for 1 h prior to IR. Cells were fixed and stained 1 h PIR. Foci intensity analysis was performed by a custom ImageJ macro. γH2AX foci were thresholded and the MFI within foci areas in the H3K27me3 channel was recorded. Subsequently, data was divided with respect to the DAPI intensity within foci area. DAPI-High indicates regions with a DAPI intensity greater than the cell-wide mean. Significance was determined by a Wilcox Ranked-Sum Test against DMSO treatment. Significance values are as follows: ns p>0.05; * p<0.05; ** p<0.01; *** p<0.001; **** p<0.0001. Three biological replicates were collected. Total number of datapoints are as follows: 1342, 1644 per group from left to right. i) Plot of the size of γH2AX foci in drug-treated MCF7 cells. Cells were treated with the indicated drugs for 60 minutes prior to dosing with 6 Gy. Cells were fixed and stained 1 h PIR. Images were acquired using a 40 X oil objective on a spinning-disk confocal microscope. Size of individual γH2AX foci were determined using a custom ImageJ macro. Significance was determined by a Wilcox Ranked-Sum Test against DMSO treatment. Significance values are as follows: ns p>0.05; * p<0.05; ** p<0.01; *** p<0.001; **** p<0.0001. Three biological replicates were collected. Total number of datapoints are as follows: 518, 516, 513, 505, 520 per group from left to right.

To further examine H3K27me3 staining after DNA damage, we applied ground state depletion (GSD) superresolution immunofluorescence imaging at 1 h PIR, revealing punctate colocalization of H3K27me3 and γH2AX staining to a 50 nm resolution (Fig. 4c). In order to directly assay molecular colocalization, we adapted GSD to enable detection of molecular proximity by Förster resonance energy transfer (FRET). Here, γH2AX was detected with an anti-γH2AX antibody coupled to a secondary antibody labeled with the donor fluorophore (DNR) and H3K27me3 with an anti-H3K27me3 antibody coupled to a secondary antibody labeled with the acceptor fluorophore (ACC). In areas where γH2AX and H3K27me3 are in molecular proximity, DNR excitation can be transferred to ACC via FRET, quenching DNR fluorescence. Imaging γH2AX and H3K27me3 at 1 h PIR in the DNR and ACC channels revealed similar distributions (Fig. 4d). Upon depletion of the ACC fluorophore by intense laser power, the H3K27me3 ACC signal was lost but the γH2AX DNR signal brightened, indicating relief of FRET quenching and thus, colocalization (Fig. 4e). A pseudocolored image indicating fold increase in DNR fluorescence after ACC depletion reveals puncta of FRET signal, consistent with H3K27me3 and γH2AX forming in molecular proximity at DSBs (Fig. 4f). Taken together, these data suggest that H3K27me3 is deposited at DSB loci and histone modifications may delineate a domain surrounding DSBs to promote detection, signaling and repair.

That a modification linked to heterochromatinization accumulates at DSBs is difficult to reconcile with purported roles for chromatin relaxation due to remodeling, PARylation and/or acetylation in γH2AX foci formation and DSB repair^31,35,89–92^. To examine whether DSB-associated H3K27me3 induces chromatin compaction, cells were stained for H3K27me3 and total H3 at 1 h PIR. H3K27me3 foci could be clearly distinguished, most of which did not appear to be associated with structures in the H3 image (Fig. 4g). Quantitation of the relative intensity of H3K27me3 and H3 staining indicated that H3K27me3 foci did not induce corresponding H3 foci (Fig. 4g right panel), arguing against local compaction and confirming focal deposition of H3K27me3 after irradiation. Proteomic data indicated that the increase in H3K27me3 was restricted to the H3.3 isoform (Fig 2d). To confirm that deposition of H3K27me3 was restricted to DSBs arising in euchromatin, we measured DAPI intensity underneath γH2AX foci as a proxy for chromatin condensation. Foci in areas with low DAPI had higher H3K27me3 levels despite, presumably, a lower density of nucleosomes in these regions (Fig 4h). Thus, we concluded that deposition of repressive chromatin marks is necessary for repair of a subset of DSBs arising in euchromatin, perhaps to attenuate local transcription. Notably, some have hypothesized that heterochromatin may be refractive to DSB induction underscoring the importance of repairing euchromatic DSBs^36,93^.

Phosphorylation of H2AX by ATM spreads kilobases away from damage sites, amplifying local signals to globally induce the DDR even from single DSBs. Thus, we assessed whether H3K27me3 might impact γH2AX spreading. Comparing the size of γH2AX foci at 1 h PIR in cells treated with vehicle, PARP inhibitor veliparib, JMJD2/4 inhibitor GSKJ4, EZH2 inhibitor GSK126 or the combination of GSK126 and GSKJ4 revealed that deregulation of H3K27me3 could impact γH2AX spreading (Fig. 4i). Strikingly, inhibition of the repressive mark H3K27me3 significantly reduced the spread of γH2AX, suggesting that PRC2 plays a role upstream of PIKKs in promoting signaling. These effects may be due to diminished H3K27me3 dependent recruitment of PRC1, a known mediator of the DDR^94^. Thus, local chromatin modification may affect global DDR signaling, and ultimately, response to IR as evidenced by radiosentization induced by EZH2 inhibitors.

## Discussion

Repair of double strand breaks is a complex process which occurs at several kinetically and spatially distinct levels in the cell. Much is known about the signaling-level events following DSB recognition (cell cycle arrest, transcriptional changes) and the downstream consequences of failure to repair DNA damage (apoptosis, senescence). However, chromatin-level changes in histone modifications which direct recognition and repair of DSBs are understudied. Indeed, DSBs are, by definition, a chromatin-localized event. Here, we report a global survey of changes to histone PTMs following DNA damage induced via ionizing radiation.

Analysis of histone PTMs after IR insult was carried out by AMT based MS/MS analysis and revealed widespread changes to the epigenome which persisted up to 48 h after IR. Modifications across all major histones were altered including modifications known to be key mediators of cell development such as H4 acetylation and H3K4 methylation. Clustering of PTM trajectories suggested at least two kinetically separate patterns of histone PTM alteration, one rapid and one occurring over ~24 hours. We chose to focus on PTMs altered at 1 h PIR as later-occurring changes are increasingly likely to mediated by cell cycle stoppage or transcriptional alteration following IR. Our analysis recapitulated several PTMs previously linked to DNA repair including H3K79me2, H3K27me3 and acetylation of the H4 tail. By assessing non-canonical PTMs via targeted PTM search for other known histone modifications we expanded the repertoire of DDR associated PTMs to include 17 types of modifications (across 3076 sites). Furthermore, though we detected small fold changes for many PTMs in our study we believe this to be reflective of larger, DSB-proximal changes diluted out by whole-chromatin analysis. Enrichment of DSB-proximal chromatin could be used to confirm our findings and definitively segregate local PTM alterations from global changes after irradiation.

Changes in other PTMs notwithstanding, we focused on alterations of H3K27 methylation and their relationship to DNA damage repair. Using MRM targeted analysis, we detected increased H3K27me3 levels following IR. We are not the first to suggest that H3K27 methylation impacts repair of DNA damage; others have presented conflicting evidence as to whether H3K27me3 or its writer, PRC2, are localized to DSBs^44,45,95^. However, to our knowledge, we are the first to use FRET imaging to localize H3K27me3 deposited on DSB proximal nucleosomes. We further suggest that H3K27me3 is a critical regulator of the DDR. Inhibition of the H3K27 methyltransferase EZH2 or the opposing demethylase, JMJD2, sensitized cells to radiation via distinct mechanisms. Blocking H3K27me3 deposition delayed break repair, while inhibiting the removal of K27 methylation precluded attenuation of DDR signaling, leading to senescence. Thus, inhibitors of H3K27 methylation are putative radiosensitizers warranting further study perhaps in an *in vivo* setting.

Towards a mechanism for H3K27 methylation in the DDR, we examined γH2AX foci establishment in the presence of H3K27 methylation inhibitors. Inhibition of either EZH2 or its counterpart JMJD2 attenuated γH2AX foci number and intensity shortly after IR insult. These data place histone modification upstream of DSB recognition by PIKKs, key mediators of downstream DDR signaling. While H3K27 trimethylation is sometimes associated with heterochromatin, EZH2 has been linked to facultative repression of genes even in non-condensed chromatin. This is in line with work which suggests that EZH2 may function to repress transcription proximal to DSB loci, thus preventing transcription across broken DNA. In our hands, we observed accumulation of H3K27me3 surrounding DSB loci without a concomitant increase histone occupancy. Thus, at early time points, DSB proximal chromatin compaction may not occur despite deposition of repressive marks. Perhaps H3K27me3 is a permissive mark which defines the DSB repair domain, or it may be required for deposition of γH2AX possibly via recruitment of ATM or another repair factors.

A role for EZH2 in preventing transcription-damage conflicts suggests that EZH2 may be specifically deposited in genic regions which sustain DNA damage. Consistent with this hypothesis, isoform-selective methylation of H3K27 following IR was observed by targeted proteomics. H3.3, an H3 isoform associated with euchromatin, realized the bulk of the increase in H3K27me3. Further, via imaging, we observed relatively stronger induction of H3K27me3 at DSBs in areas of open chromatin, likely euchromatin. Additionally, we place H3K27me3 upstream of R-Loop formation and show that inhibiting EZH2 prevents attenuation of Pol II at DSBs. Collectively, these data raise the possibility that different genomic regions may require distinct repair programs dependent upon their basal epigenetic state. By extension, the role of histone marks in directing DSB repair may be distinct from their basal location or activity.

It is interesting, and indeed apparently paradoxical, that either increased or decreased DSB-proximal H3K27me3 levels are sufficient to attenuate γH2AX deposition. However, we note that inhibition of EZH2 or JMJD2 evinced different phenotypes, with only the latter accelerating cellular senescence. Additionally, we posit that our findings could be evidence of a multistep process of histone methylation at DSBs which is separated either kinetically or spatially. For example, it may be that H3K27me3 deposition is necessary for repair in euchromatin, but this excess methylation must later be removed to restore basal chromatin activity. Failure to restore the basal epigenetic state may prolong DDR signaling and contribute to senescence. Indeed, altering H3K27me3 levels has been shown to induce senescence in the absence of DNA damage^96^. Our data is also consistent with reports that the H3 demethylase UTX is required for the DDR^97^. Returning to the influence of basal epigenetic states on the DDR, loci in different epigenetic states may be repaired via distinct epigenetic mechanisms or at different times following IR. This is consistent with the separable kinetics of histone modifications observed in our proteomics data. Future studies must address the relationship between preexisting chromatin state, repair pathway and repair kinetics. For example, many studies note special repair pathways and activities for heterochromatic regions or telomeric chromatin^30,98^.

Our findings also suggest a more fundamental purpose of highly conserved epigenetic readers and writers such as the polycomb family. PRC2 was first identified in flies and is highly conserved even in organisms which lack complex gene expression control^99,100^. Yet, all eukaryotes have DNA repair systems to repair breaks and safeguard genetic information. Therefore, it is likely that the DDR activity of PRC2 and other enzymes does not represent moonlighting, but rather is an essential and ancient subset of their functions. A fuller understanding of how these enzymes function in DSB repair may, in turn, shed light on their roles in transcription. Transcription-coupled repair of DSBs has been postulated, as has transcriptional damage to DNA^101,102^. This study reframes these concepts by suggesting transcription-independent roles for transcriptional machinery in the DDR.

## Abbreviations

AMT: Accurate Mass and time
ATM: Ataxia telangiectasia mutated
ATR: Ataxia telangiectasia and Rad3 related
DDR: DNA Damage Response
DSB: Double Strand Break
DNA-PKcs: DNA-dependent Protein Kinase
EHMP: Epiproteomic Histone Modification Panel
EZH2: Enhancer of Zest Homologue 2
FRET: Förster Resonance Energy Transfer
γH2AX: gamma H2AX is the phosphorylated form of H2AX
HR: Homologous Recombination
IR: Ionizing Radiation
IRIF: Ionizing Radiation Induced Foci
NHEJ: Non-homologous End Joining
MRM: Multiple Reaction Monitoring
PARP: Poly ADP-ribose Polymerase
PIKK: Phosphatidylinositol 3-kinase-related kinases
PIR: Post-IR
Pol II: RNA Polymerase II
PRC2: Polycomb Repressive Complex 2
PTM: Post Translational Modification

## Data Availability

All data and code used to generate figures are available upon request to the corresponding author. Epiprofile proteomics data have been deposited to the ProteomeXchange Consortium via the PRIDE partner repository^62^ with the dataset identifier PXD019388. EHMP data is attached to this manuscript.

## Acknowledgements

We thank Jacek Sikora (now at AbbVie)and the staff of Northwestern Proteomics and Ken Johnson and the staff of the Mayo Clinic Proteomics Core for proteomics support and Vytas Bindokas and Christine Labno of the University of Chicago Integrated Light Microscopy Core for training and consultation.

## Grant Funding

This work was supported by NCI grants R21 CA213247 and R01 CA199663 and by DoD CDMRP PRCRP Impact Award CA190982 to SJK. JL was partially supported by the Multi-disciplinary Training program in Cancer Research (MTCR), T32 CA009594. The Integrated Light Microscopy Core was supported by NCI cancer center grant P30 CA014599. Northwestern Proteomics was supported by P30 CA060553 and P41 GM108569.

## Contributions

J.L. conceived the experiments, performed experiments, obtained and analyzed images, assembled the data and wrote the manuscript. D.W. prepared proteomic samples, analyzed LC-MS/MS data and helped prepare the manuscript. S.K. supervised the study. All authors read and approved the final manuscript.

## Ethics declarations

The authors declare no competing interests

## Supplemental Data

This article contains supplemental data.

## Supplemental Figures

**Figure S1 EHMP analysis reveals several Histone PTMs are altered following IR**

a) Select H3 and H4 residue PTM data from the EHMP survey are shown. Height of the bars represents the percent of the residue modified at the indicated timepoint. Error bars show standard deviation for three replicates. Significance was determined by a Kruskall-Wallace test comparing timepoints within a given residue. Significance values are as follows: ns p>0.05; * p<0.05; ** p<0.01; *** p<0.001; **** p<0.0001.

**Figure S2 EpiProfiler confirms IR mediated changes to histone PTMs**

a) Select H3 and H4 residue PTM data from the EpiProfiler Histone PTM dataset are shown. Height of the bars represents the percent of the residue modified at the indicated timepoint. Error bars show standard deviation for three replicates. Significance was determined by a Friedman test comparing timepoints within a given residue. Significance values are as follows: ns p>0.05; * p<0.05; ** p<0.01; *** p<0.001; **** p<0.0001.

**Figure S3 EHMP and Epiprofile analyses report differential histone alterations**

a) Matrix shows the correlation between 55 PTMs measured by both EpiProfile and the EHMP assay. Color of the squares is proportional to Pearsons correlation coefficient. The R^2^ value between two timepoints is shown within each square. Data used to construct the matrix are the average percent residue modification values.

**Figure S4 Inhibition of H3K27 methylation sensitizes cells to Ionizing Radiation**

a) H3K27me3 mean fluorescent intensity of drug treated cells. Cells were treated and imaged as in Fig. 3b. MFI is calculated for each nucleus using a custom ImageJ macro. Significance was determined by a Wilcox Ranked-Sum Test against DMSO treatment. Significance values are as follows: ns p>0.05; * p<0.05; ** p<0.01; *** p<0.001; **** p<0.0001. Three biological replicates were collected. Total number of datapoints are as follows: 110, 62, 43, 49, 38 per group from left to right.

b) Incucyte growth curves of drug treated cells. Cells were treated for 60 min as in Fig. 3b and then exposed to IR or mock irradiated (NIR). Cell number was tracked for 120 h in an Incucyte system. The mean normalized number of cells is plotted, and error bars denote SEM for 3 replicates. Significance was determined by Dunnett’s Multiple Comparisons Test against DMSO treatment.

c) Images excerpted from the Incucyte image dataset over the course of the 120 h analysis. Timepoints are equivalent to Fig. 3d. Only the IR condition is shown.

d) SA-βGal staining of cells treated for 1 h with the indicated drugs prior to IR insult and allowed to recover for 72 h before fixation and staining. A representative image, selected from three replicates, is shown for each treatment.

## Notes

### Competing Interest Statement

The authors have declared no competing interest.

## References

1. Pilié, P. G., Tang, C., Mills, G. B. and Yap, T. A. State-of-the-art strategies for targeting the DNA damage response in cancer. Nature Reviews Clinical Oncology 16, 81–104 (2018)

2. Lomax, M. E., Folkes, L. K. and O’Neill, P. Biological Consequences of Radiation-induced DNA Damage: Relevance to Radiotherapy. Clinical Oncology 25, 578–585 (2013).

3. Orth, M., Lauber, K., Niyazi, M., Friedl, A. A., Li, M., Maihöfer, C., Schüttrumpf, L., Ernst, A., Niemöller, O. M. and Belka, C. Current concepts in clinical radiation oncology. Radiation and Environmental Biophysics 53, 1–29 (2014).

4. Xiao, Y. and Rosen, M. The role of Imaging and Radiation Oncology Core for precision medicine era of clinical trial. Translational Lung Cancer Research 6, 621–624 (2017).

5. Conibear, J. Rationale for concurrent chemoradiotherapy for patients with stage III non-small-cell lung cancer. British Journal of Cancer 123, 10–17 (2020)

6. Machtay, M., Moughan, J., Trotti, A., Garden, A. S., Weber, R. S., Cooper, J. S., Forastiere, A. and Ang, K. K. Factors Associated With Severe Late Toxicity After Concurrent Chemoradiation for Locally Advanced Head and Neck Cancer: An RTOG Analysis. Journal of Clinical Oncology 26, (2021).

7. Du, C., Ying, H., Kong, F., Zhai, R. and Hu, C. Concurrent chemoradiotherapy was associated with a higher severe late toxicity rate in nasopharyngeal carcinoma patients compared with radiotherapy alone: a meta-analysis based on randomized controlled trials. Radiation Oncology 10, 70 (2015).

8. Curtin, N. J. DNA repair dysregulation from cancer driver to therapeutic target. Nature Reviews Cancer 12, 801–817 (2012).

9. De Schutter, H. and Nuyts, S. Radiosensitizing potential of epigenetic anticancer drugs. Anti-cancer agents in medicinal chemistry 9, 99–108 (2009).

10. Jachimowicz, R. D., Goergens, J. and Reinhardt, H. C. DNA double-strand break repair pathway choice - from basic biology to clinical exploitation. Cell Cycle vol. 18 1423–1434 (2019).

11. Sonnenblick, A., De Azambuja, E., Azim, H. A. and Piccart, M. An update on PARP inhibitors—moving to the adjuvant setting. Nature Publishing Group, 12, 27–41 (2014).

12. D’Andrea, A. D. Mechanisms of PARP inhibitor sensitivity and resistance. DNA Repair 71:172–176 (2018).

13. Clouaire, T. and Legube, G. A Snapshot on the Cis Chromatin Response to DNA Double-Strand Breaks. Trends in Genetics 35, 330–345 (2019).

14. Yamamori, T., Yasui, H., Yamazumi, M., Wada, Y., Nakamura, Y., Nakamura, H. and Inanami, O. Ionizing radiation induces mitochondrial reactive oxygen species production accompanied by upregulation of mitochondrial electron transport chain function and mitochondrial content under control of the cell cycle checkpoint. Free Radical Biology and Medicine 53, 260–270 (2012).

15. Rogakou, E. P., Pilch, D. R., Orr, A. H., Ivanova, V. S. and Bonner, W. M. DNA double-stranded breaks induce histone H2AX phosphorylation on serine 139. The Journal of biological chemistry 273, 5858–68 (1998).

16. Paull, T. T., Rogakou, E. P., Yamazaki, V., Kirchgessner, C. U., Gellert, M. and Bonner, W. M. A critical role for histone H2AX in recruitment of repair factors to nuclear foci after DNA damage. Current Biology 10, 886–895 (2000).

17. Rothkamm, K., Barnard, S., Moquet, J., Ellender, M., Rana, Z. and Burdak‐Rothkamm, S. DNA damage foci: Meaning and significance. Environmental and molecular mutagenesis 56, 491–504 (2015).

18. Belyaev, I. Y. Radiation-induced DNA repair foci: spatio-temporal aspects of formation, application for assessment of radiosensitivity and biological dosimetry. Mutation Research/Reviews in Mutation Research 704, 132–141 (2010).

19. Bai, P. Biology of Poly(ADP-Ribose) Polymerases: The Factotums of Cell Maintenance. Molecular Cell 58, 947–958 (2015).

20. Gupte, R., Liu, Z. and Kraus, W. L. PARPs and ADP-ribosylation: Recent advances linking molecular functions to biological outcomes. Genes and Development 31, 101–126 (2017).

21. Skidmore, C. J., Davies, M. I., Goodwin, P. M., Halldorsson, H., Lewis, P. J., Shall, S. and Zia’ee, A. A. The Involvement of Poly(ADP‐ribose) Polymerase in the Degradation of NAD Caused by γ‐Radiation and N‐Methyl‐N‐Nitrosourea. European Journal of Biochemistry 101, 135–142 (1979).

22. Chapman, J. R., Taylor, M. R. G. and Boulton, S. J. Playing the End Game: DNA Double-Strand Break Repair Pathway Choice. Molecular Cell 47, 497–510 (2012).

23. Ceccaldi, R., Rondinelli, B. and D’Andrea, A. D. Repair Pathway Choices and Consequences at the Double-Strand Break. Trends in Cell Biology vol. 26 52–64 (2016).

24. Panier, S. and Boulton, S. J. Double-strand break repair: 53BP1 comes into focus. Nature Reviews Molecular Cell Biology 15, 7–18 (2013).

25. Jackson, S. P. and Bartek, J. The DNA-damage response in human biology and disease. Nature 461, 1071–1078 (2009).

26. Blackford, A. N. and Jackson, S. P. Molecular Cell Review ATM, ATR, and DNA-PK: The Trinity at the Heart of the DNA Damage Response. Molecular Cell 66, 801–817 (2017).

27. Arnoult, N., Correia, A., Ma, J., Merlo, A., Garcia-Gomez, S., Maric, M., Tognetti, M., Benner, C. W., Boulton, S.J., Saghatelian, A. and Karlseder, J. Regulation of DNA repair pathway choice in S and G2 phases by the NHEJ inhibitor CYREN. Nature 549, 548–552 (2017).

28. Jeggo, P. A., Downs, J. A. and Gasser, S. M. Chromatin modifiers and remodelers in DNA repair and signaling. Philosophical Transactions of the Royal Society B: Biological Sciences 372, 20160279 (2017).

29. Hunt, Clayton R., Deepti Ramnarain, Nobuo Horikoshi, Puneeth Iyengar, Raj K. Pandita, Jerry W. Shay, and Tej K. Pandita. Histone Modifications and DNA Double-Strand Break Repair after Exposure to Ionizing Radiations. Radiation Research 179, 383–392. 2013.

30. Chiolo, I., Caridi, P. C., Delabaere, L. and Zapotoczny, G. And yet, it moves: nuclear and chromatin dynamics of a heterochromatic double-strand break. Philosophical transactions of the Royal Society of London. Series B, Biological sciences 372, (2017).

31. Burgess, R. C., Burman, B., Kruhlak, M. J. and Misteli, T. Activation of DNA Damage Response Signaling by Condensed Chromatin. Cell Reports 9, 1703–1718 (2014).

32. Delgoffe, Greg M., Kristen N. Pollizzi, Adam T. Waickman, Emily Heikamp, David J. Meyers, Maureen R. Horton, Bo Xiao, Paul F. Worley, Jonathan D. Powell. The kinase mTOR regulates the differentiation of helper T cells through the selective activation of signaling by mTORC1 and mTORC2. Nature immunology 12, 295–303 (2011).

33. Ségurel, L. and Bon, C. Recent Advancements in DNA Damage–Transcription Crosstalk and High-Resolution Mapping of DNA Breaks. Annu. Rev. Genome Human Genetics 18, 87–113 (2017).

34. Lavelle, C. and Foray, N. Chromatin structure and radiation-induced DNA damage: From structural biology to radiobiology. International Journal of Biochemistry and Cell Biology 49, 84–97 (2014).

35. Dellaire, G., Kepkay, R. and Bazett-Jones, D. P. High resolution imaging of changes in the structure and spatial organization of chromatin, γ-H2A.X and the MRN complex within etoposide-induced DNA repair foci. Cell Cycle 8, 3750–3769 (2009).

36. Kim, J. A., Kruhlak, M., Dotiwala, F., Nussenzweig, A. and Haber, J. E. Heterochromatin is refractory to γ-H2AX modification in yeast and mammals. Journal of Cell Biology 178, 209–218 (2007).

37. Murga, M., Jaco, I., Fan, Y., Soria, R., Martinez-Pastor, B., Cuadrado, M., Yang, S.M., Blasco, M. A., Skoultchi, A. I. and Fernandez-Capetillo, O. Global chromatin compaction limits the strength of the DNA damage response. Journal of Cell Biology 178, 1101–1108 (2007).

38. Takata, H., Hanafusa, T., Mori, T., Shimura, M., Iida, Y., Ishikawa, K., Yoshikawa, K., Yoshikawa, Y. and Maeshima, K. Chromatin Compaction Protects Genomic DNA from Radiation Damage. PLoS ONE 8, 1–11 (2013).

39. Spotheim-Maurizot, M., Ruiz, S., Sabattier, R. and Charlier, M. Radioprotection of DNA by Polyamines. International Journal of Radiation Biology 68, 571–577 (1995).

40. Schuettengruber, B., Bourbon, H. M., Di Croce, L. and Cavalli, G. Leading Edge Review Genome Regulation by Polycomb and Trithorax: 70 Years and Counting. Cell 171, 34–57 (2017).

41. van Kruijsbergen, I., Hontelez, S. and Veenstra, G. J. C. Recruiting polycomb to chromatin. International Journal of Biochemistry and Cell Biology 67, 177–187 (2015).

42. di Croce, L. and Helin, K. Transcriptional regulation by Polycomb group proteins. Nature Structural and Molecular Biology 20, 1147–1155 (2013).

43. Aranda, S., Mas, G. and di Croce, L. Regulation of gene transcription by Polycomb proteins. Science Advances 1, 1–16 (2015).

44. Campbell, S., Ismail, I. H., Young, L. C., Poirier, G. G. and Hendzel, M. J. Polycomb repressive complex 2 contributes to DNA double-strand break repair. Cell Cycle 12, 2675–2683 (2013).

45. Chou, D. M., Adamson, B., Dephoure, N. E., Tan, X., Nottke, A. C., Hurov, K. E., Gygi, S. P., Colaiácovo, M. P. and Elledge, S. J. A chromatin localization screen reveals poly (ADP ribose)-regulated recruitment of the repressive polycomb and NuRD complexes to sites of DNA damage. Proceedings of the National Academy of Sciences 107, 18475–18480 (2010).

46. Clouaire, T., Rocher, V., Lashgari, A., Arnould, C., Aguirrebengoa, M., Biernacka, A., Skrzypczak, M., Aymard, F., Fongang, B., Dojer, N. and Iacovoni, J.S. Comprehensive Mapping of Histone Modifications at DNA Double-Strand Breaks Deciphers Repair Pathway Chromatin Signatures. Molecular Cell 72, 250–262. (2018).

47. Piunti, A. and Shilatifard, A. The roles of Polycomb repressive complexes in mammalian development and cancer. Nature Reviews Molecular Cell Biology 22, 326–345 (2021).

48. Zhang Y., Chang J. F., Sun J., Chen L., Yang X. M., Tang H. Y., Jing Y. Y., Kang X, He Z. M., Wu, J. Y., and Wei, H. M. Histone H3K27 methylation is required for NHEJ and genome stability by modulating the dynamics of FANCD2 on chromatin. Journal of cell science 131.12 (2018).

49. Efimova E. V, Takahashi S, Shamsi N. A., Wu D., Labay E, Ulanovskaya OA, Weichselbaum RR, Kozmin SA, Kron SJ. DNA Damage and Repair Linking Cancer Metabolism to DNA Repair and Accelerated Senescence. Mol Cancer Res 173–184 (2016)

50. Caruso L. B., Martin K. A., Lauretti E., Hulse M., Siciliano M., Lupey-Green L.N., Abraham A., Skorski T., Tempera I. Poly(ADP-ribose) Polymerase 1, PARP1, modifies EZH2 and inhibits EZH2 histone methyltransferase activity after DNA damage. Oncotarget 9, 10585–10605 (2018).

51. Li J, Hart R. P., Mallimo E. M., Swerdel M. R., Kusnecov, A. W., Herrup K. EZH2-mediated H3K27 trimethylation mediates neurodegeneration in ataxia-telangiectasia. Nature Neuroscience 16, 1745–53. (2013).

52. Wang Y., Sun H., Wang J., Wang H., Meng L., Xu C., Jin M., Wang B., Zhang Y., Zhu T. DNA-PK-mediated phosphorylation of EZH2 regulates the DNA damage-induced apoptosis to maintain T-cell genomic integrity. Cell Death and Disease 7, 1–10 (2016).

53. Finlay, M. R. V. and Griffin, R. J. Modulation of DNA repair by pharmacological inhibitors of the PIKK protein kinase family. Bioorganic and Medicinal Chemistry Letters 22, 5352–5359 (2012).

54. Bryant H. E., Schultz N., Thomas H. D., Parker K. M., Flower D., Lopez E., Kyle S., Meuth M., Curtin N. J., Helleday, T. Specific killing of BRCA2-deficient tumours with inhibitors of poly(ADP-ribose) polymerase. Nature 434, 913–917 (2005).

55. Garcia B. A., Mollah S., Ueberheide B. M., Busby S. A., Muratore T. L., Shabanowitz J., Hunt D. F. Chemical derivatization of histones for facilitated analysis by mass spectrometry. Nature Protocols 2, 933–938 (2007).

56. MacLean B., Tomazela D. M., Shulman N., Chambers M., Finney G. L., Frewen B., Kern R., Tabb D. L., Liebler D. C., MacCoss M. J. Skyline: An open source document editor for creating and analyzing targeted proteomics experiments. Bioinformatics 26, 966–968 (2010).

57. Yuan Z. F., Liu C., Wang H. P., Sun R. X., Fu Y., Zhang J. F., Wang L. H., Chi H., Li Y., Xiu L. Y., Wang W. P. pParse: A method for accurate determination of monoisotopic peaks in high-resolution mass spectra. Proteomics 12, 226–235 (2012).

58. Yuan Z. F., Sidoli S., Marchione D. M., Simithy J., Janssen K. A., Szurgot M. R., Garcia B. A. EpiProfile 2.0: A Computational Platform for Processing Epi-Proteomics Mass Spectrometry Data. Journal of Proteome Research 17, 2533–2541 (2018).

59. Yuan Z. F, Lin S., Molden R. C., Cao X. J., Bhanu N. V., Wang X., Sidoli S., Liu S., Garcia B. A. EpiProfile Quantifies Histone Peptides with Modifications by Extracting Retention Time and Intensity in High-resolution Mass Spectra. Molecular and Cellular Proteomics 14, 1696–1707 (2015).

60. Tyanova S., Temu T., Sinitcyn P., Carlson A., Hein M. Y., Geiger T., Mann M., Cox J. The Perseus computational platform for comprehensive analysis of (prote)omics data. Nature Methods 13, 731–740 (2016).

61. Deutsch E. W., Bandeira N., Sharma V., Perez-Riverol Y., Carver J. J., Kundu D. J, García-Seisdedos D., Jarnuczak A. F., Hewapathirana S., Pullman B. S., Wertz J. The ProteomeXchange consortium in 2020: Enabling “big data” approaches in proteomics. Nucleic Acids Research 48, D1145–D1152 (2020).

62. Perez-Riverol Y., Csordas A., Bai J., Bernal-Llinares M., Hewapathirana S., Kundu D. J., Inuganti A., Griss J., Mayer G., Eisenacher M., Pérez E. The PRIDE database and related tools and resources in 2019: Improving support for quantification data. Nucleic Acids Research 47, D442–D450 (2019).

63. Gyori, B. M., Venkatachalam, G., Thiagarajan, P. S., Hsu, D. and Clement, M. V. OpenComet: An automated tool for comet assay image analysis. Redox Biology 2, 457–465 (2014).

64. Bolte, S. and Cordelières, F. P. A guided tour into subcellular colocalization analysis in light microscopy. Journal of Microscopy 224, 213–232 (2006).

65. Ovesný, M., Křížek, P., Borkovec, J., Švindrych, Z. and Hagen, G. M. ThunderSTORM: a comprehensive ImageJ plug-in for PALM and STORM data analysis and super-resolution imaging. Bioinformatics 30, 2389–2390 (2014).

66. Kim, J. J., Lee, S. Y., and Miller, K. M. Preserving genome integrity and function: the DNA damage response and histone modifications. Critical Reviews in Biochemistry and Molecular Biology 54, 208–241 (2019).

67. Polo, S. E. and Jackson, S. P. Dynamics of DNA damage response proteins at DNA breaks: A focus on protein modifications. Genes and Development 25, 409–433 (2011).

68. von Stechow, L. and Olsen, J. V. Proteomics insights into DNA damage response and translating this knowledge to clinical strategies. Proteomics 17, 1600018 (2017).

69. Arnaudo, A. M. and Garcia, B. A. Proteomic characterization of novel histone post-translational modifications. Epigenetics and Chromatin 6, 1–7 (2013).

70. Huang, H., Lin, S., Garcia, B. A. and Zhao, Y. Quantitative proteomic analysis of histone modifications. Chemical Reviews 115, 2376–2418 (2015).

71. Sudprasert, W., Navasumrit, P. and Ruchirawat, M. Effects of low-dose gamma radiation on DNA damage, chromosomal aberration and expression of repair genes in human blood cells. International Journal of Hygiene and Environmental Health 209, 503–511 (2006).

72. Banáth, J. P., MacPhail, S. H. and Olive, P. L. Radiation sensitivity, H2AX phosphorylation, and kinetics of repair of DNA strand breaks in irradiated cervical cancer cell lines. Cancer Research 64, 7144–7149 (2004).

73. Dhar, S., Gursoy-yuzugullu, O., Parasuram, R., and Price, B. D. The tale of a tail: histone H4 acetylation and the repair of DNA breaks. Philosophical Transactions of the Royal Society B: Biological Sciences 372, 20160284 (2017).

74. Nguyen, A. T. and Zhang, Y. The diverse functions of Dot1 and H3K79 methylation. Genes and development 25, 1345–58 (2011).

75. Goldberg, A. D., Banaszynski, L. A., Noh, K. M., Lewis, P. W., Elsaesser, S. J., Stadler, S., Dewell, S., Law, M., Guo, X., Li, X. and Wen, D. Distinct Factors Control Histone Variant H3.3 Localization at Specific Genomic Regions. Cell 140, 678–691 (2010).

76. Boros, J., Arnoult, N., Stroobant, V., Collet, J. F. and Decottignies, A. Polycomb repressive complex 2 and H3K27me3 cooperate with H3K9 methylation to maintain heterochromatin protein 1α at chromatin. Molecular and cellular biology 34, 3662–74 (2014).

77. Wiles, E. T. and Selker, E. U. H3K27 methylation: a promiscuous repressive chromatin mark. Current Opinion in Genetics and Development 43, 31–37 (2017).

78. Izhar, L., Adamson, B., Ciccia, A., Lewis, J., Pontano-Vaites, L., Leng, Y., Liang, A.C., Westbrook, T.F., Harper, J.W. and Elledge, S.J. A Systematic Analysis of Factors Localized to Damaged Chromatin Reveals PARP-Dependent Recruitment of Transcription Factors. CellReports 11, 1486–1500 (2015).

79. Nichol, J. N., Dupéré-Richer, D., Ezponda, T., Licht, J. D. and Miller, W. H. H3K27 Methylation: A Focal Point of Epigenetic Deregulation in Cancer. Advances in Cancer Research 131, 59–95 (2016).

80. Agger, K., Cloos, P. A., Christensen, J., Pasini, D., Rose, S., Rappsilber, J., Issaeva, I., Canaani, E., Salcini, A.E. and Helin, K. UTX and JMJD3 are histone H3K27 demethylases involved in HOX gene regulation and development. Nature 449, 731–734 (2007).

81. Murai, J., Shar-yin, N. H., Das, B. B., Renaud, A., Zhang, Y., Doroshow, J. H., Ji, J., Takeda, S. and Pommier, Y. Trapping of PARP1 and PARP2 by clinical PARP inhibitors. Cancer Research 72, 5588–5599 (2012).

82. Sollier, J. and Cimprich, K. A. Breaking bad: R-loops and genome integrity. Trends in Cell Biology 25, 514–522 (2015).

83. Aguilera, A. and Gómez-González, B. DNA-RNA hybrids: The risks of DNA breakage during transcription. Nature Structural and Molecular Biology 24, 439–443 (2017).

84. Feringa, F. M., Raaijmakers, J. A., Hadders, M. A., Vaarting, C., Macurek, L., Heitink, L., Krenning, L. and Medema, R. H. Persistent repair intermediates induce senescence. Nature Communications 9, 3923 (2018).

85. Labay, E., Efimova, E. V., Quarshie, B. K., Golden, D. W., Weichselbaum, R. R. and Kron, S. J. Ionizing radiation-induced foci persistence screen to discover enhancers of accelerated senescence. International journal of high throughput screening 2, 1 (2011).

86. Fumagalli, M., Rossiello, F., Mondello, C. and D’Adda Di Fagagna, F. Stable cellular senescence is associated with persistent DDR activation. PLoS ONE 9, 44–46 (2014).

87. Liu, Y., Efimova, E. V., Ramamurthy, A. and Kron, S. J. Repair-independent functions of DNA-PKcs protect irradiated cells from mitotic slippage and accelerated senescence. Journal of Cell Science 132, 13 (2019).

88. Fumagalli, M., Rossiello, F., Clerici, M., Barozzi, S., Cittaro, D., Kaplunov, J. M., Bucci, G., Dobreva, M., Matti, V., Beausejour, C. M. and Herbig, U. Telomeric DNA damage is irreparable and causes persistent DNA-damage-response activation. Nature Cell Biology 14, 355–365 (2012).

89. Ogiwara, H., Ui, A., Otsuka, A., Satoh, H., Yokomi, I., Nakajima, S., Yasui, A., Yokota, J. and Kohno, T. Histone acetylation by CBP and p300 at double-strand break sites facilitates SWI/SNF chromatin remodeling and the recruitment of non-homologous end joining factors. Oncogene 30, 2135–2146 (2011).

90. Kruhlak, M. J., Celeste, A., Dellaire, G., Fernandez-Capetillo, O., MuLJller, W. G., McNally, J. G., Bazett-Jones, D. P. and Nussenzweig, A. Changes in chromatin structure and mobility in living cells at sites of DNA double-strand breaks. The Journal of Cell Biology 172, 823–34 (2006)

91. Ziv, Y., Bielopolski, D., Galanty, Y., Lukas, C., Taya, Y., Schultz, D. C., Lukas, J., Bekker-Jensen, S., Bartek, J. and Shiloh, Y. Chromatin relaxation in response to DNA double-strand breaks is modulated by a novel ATM-and KAP-1 dependent pathway. Nature Cell Biology 8, 870–876 (2006).

92. Sellou, H., Lebeaupin, T., Chapuis, C., Smith, R., Hegele, A., Singh, H. R., Kozlowski, M., Bultmann, S., Ladurner, A. G., Timinszky, G. and Huet, S. The poly(ADP-ribose)-dependent chromatin remodeler Alc1 induces local chromatin relaxation upon DNA damage. Molecular Biology of the Cell 27, 3791–3799 (2016).

93. Barone, F., Belli, M., Pazzaglia, S., Sapora, O. and Tabocchini, M. A. Radiation damage and chromatin structure. Annali dell’Istituto superiore di sanita 25, 59—67 (1989).

94. Vissers, J. H. A., van Lohuizen, M. and Citterio, E. The emerging role of Polycomb repressors in the response to DNA damage. Journal of Cell Science 125, 3939–3948 (2012).

95. Gong, F., Chiu, L. Y., Cox, B., Aymard, F., Clouaire, T., Leung, J. W., Cammarata, M., Perez, M., Agarwal, P., Brodbelt, J. S. and Legube, G. Screen identifies bromodomain protein ZMYND8 in chromatin recognition of transcription-associated DNA damage that promotes homologous recombination. Genes and Development 29, 197–211 (2015).

96. Ito, T., Teo, Y. V., Evans, S. A., Neretti, N. and Sedivy, J. M. Regulation of Cellular Senescence by Polycomb Chromatin Modifiers through Distinct DNA Damage- and Histone Methylation-Dependent Pathways. Cell Reports 22, 3480–3492 (2018).

97. Rath, B. H., Waung, I., Camphausen, K. and Tofilon, P. J. Inhibition of the histone h3k27 demethylase utx enhances tumor cell radiosensitivity. Molecular Cancer Therapeutics 17, 1070–1078 (2018).

98. Webb, C. J., Wu, Y. and Zakian, V. A. DNA repair at telomeres: keeping the ends intact. Cold Spring Harbor perspectives in biology 5, (2013).

99. Margueron, R. and Reinberg, D. The Polycomb complex PRC2 and its mark in life. Nature 469, 343–349 (2011).

100. Lewis, E. B. A gene complex controlling segmentation in Drosophila. Nature 276, 565–570 (1978).

101. Marnef, A., Cohen, S. and Legube, G. Transcription-Coupled DNA Double-Strand Break Repair: Active Genes Need Special Care. Journal of Molecular Biology 429, 1277–1288 (2017).

102. Gregersen, L. H. and Svejstrup, J. Q. The Cellular Response to Transcription-Blocking DNA Damage. Trends in Biochemical Sciences 43, 327–341 (2018).

