## Supplemental Figs for "Global Epigenetic Analysis Reveals H3K27 Methylation as a Mediator of Double Strand Break Repair"

Supplemental Files Table of Contents:

FigS1: EHMP analysis reveals several Histone PTMs are altered following IR
FigS2: EpiProfiler confirms IR mediated changes to histone PTMs
FigS3: EHMP and Epiprofile analyses report differential histone alterations
FigS4: Inhibition of H3K27 methylation sensitizes cells to Ionizing Radiation

Table S1: Raw EMHP Percent Residue Modification Data
Table S2: DTW Cluster Assignments
Table S3: Results of Statistical Tests for all EHMP PTMs
Table S4: Raw EpiProfiler Percent Residue Modification Data and Peptide Intensities
Table S5: Summary of Manual MaxQuant Search
Table S6: All Additional Histone PTMs Detected by Manual MaxQuant Search

a

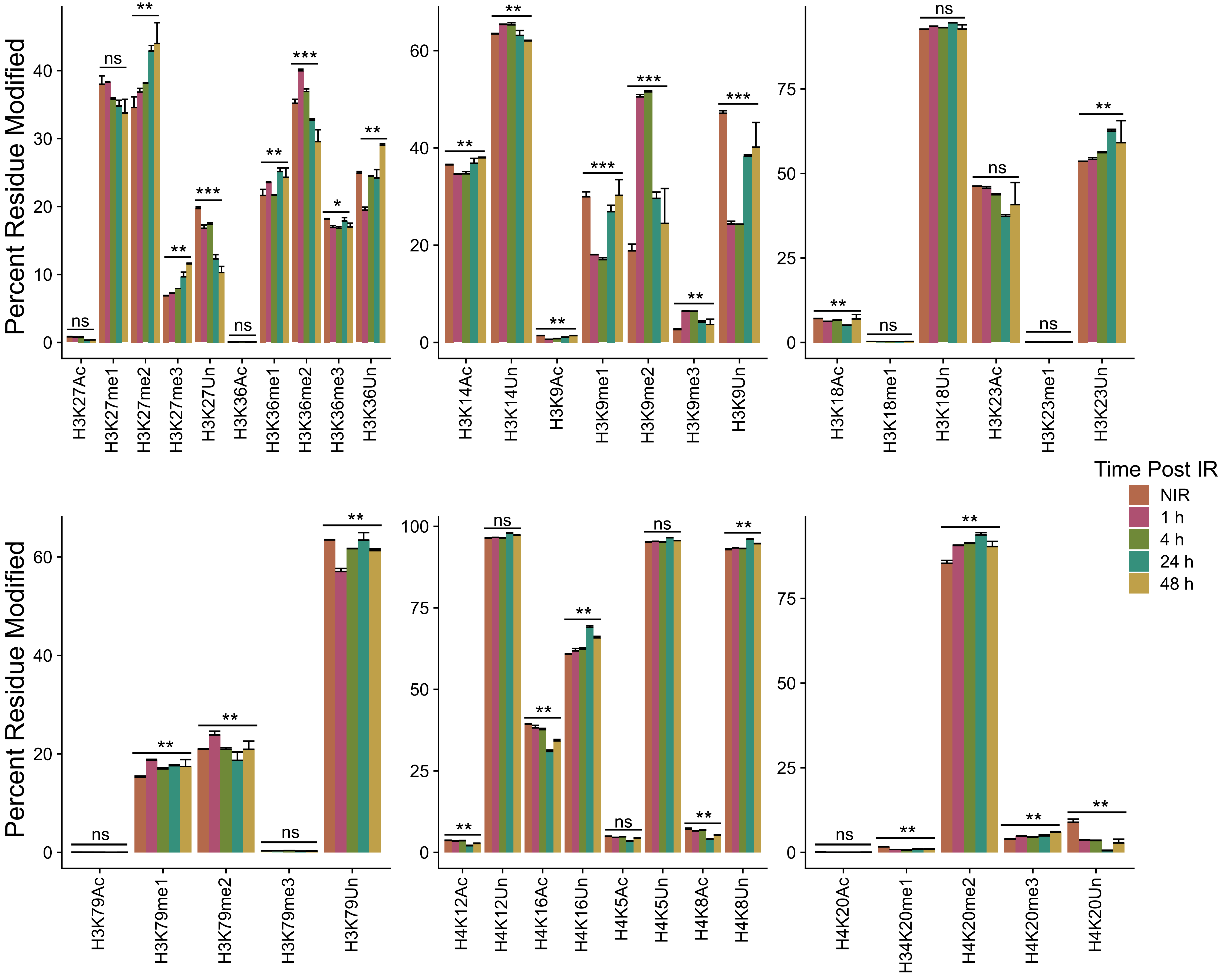
Percent Residue Modified
H3K27Ac
H3K27me1
H3K27me2
H3K27me3
H3K27Un
H3K36Ac
H3K36me1
H3K36me2
H3K36me3
H3K36Un
H3K14Ac
H3K14Un
H3K9Ac
H3K9me1
H3K9me2
H3K9me3
H3K9Un
H3K18Ac
H3K18me1
H3K18Un
H3K23Ac
H3K23me1
H3K23Un
H3K79Ac
H3K79me1
H3K79me2
H3K79me3
H3K79Un
H4K12Ac
H4K12Un
H4K16Ac
H4K16Un
H4K5Ac
H4K5Un
H4K8Ac
H4K8Un
H4K20Ac
H34K20me1
H4K20me2
H4K20me3
H4K20Un
Time Post IR
NIR
1 h
4 h
24 h
48 h

**Figure S1 EHMP analysis reveals several Histone PTMs are altered following IR**

a) Select H3 and H4 residue PTM data from the EHMP survey are shown. Height of the bars represents the percent of the residue modified at the indicated timepoint. Error bars show standard deviation for three replicates. Significance was determined by a Kruskall-Wallace test comparing timepoints within a given residue. Significance values are as follows: ns $p>0.05$; * $p<0.05$; ** $p<0.01$; *** $p<0.001$; **** $p<0.0001$.

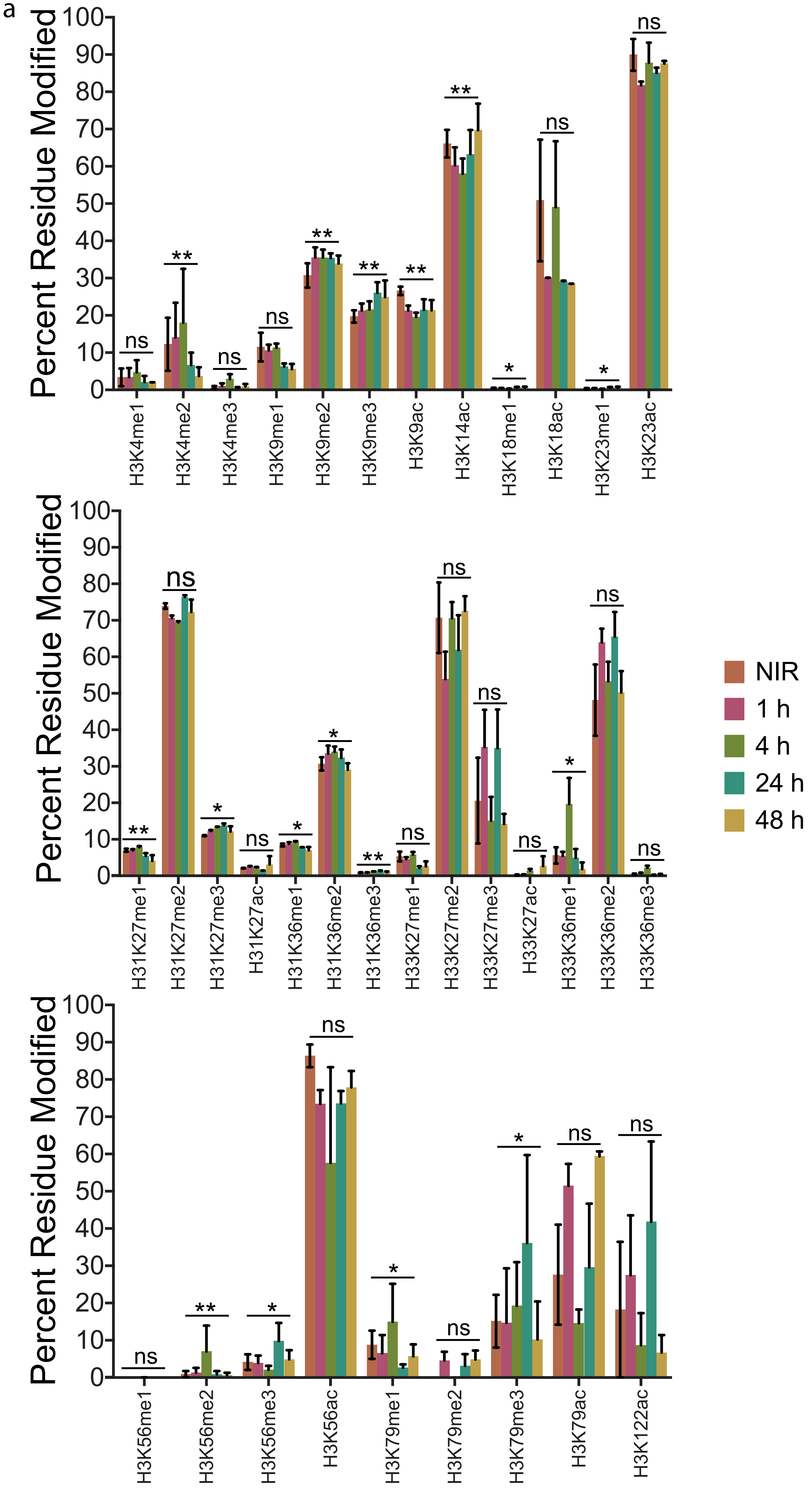

a
Percent Residue Modified
ns
**
ns
ns
**
**
**
**
*
ns
*
ns
H3K4me1
H3K4me2
H3K4me3
H3K9me1
H3K9me2
H3K9me3
H3K9ac
H3K14ac
H3K18me1
H3K18ac
H3K23me1
H3K23ac
Percent Residue Modified
**
ns
*
ns
*
*
**
ns
ns
ns
ns
*
ns
ns
H31K27me1
H31K27me2
H31K27me3
H31K27ac
H31K36me1
H31K36me2
H31K36me3
H33K27me1
H33K27me2
H33K27me3
H33K27ac
H33K36me1
H33K36me2
H33K36me3
NIR
1 h
4 h
24 h
48 h
Percent Residue Modified
ns
**
*
ns
*
ns
*
ns
ns
H3K56me1
H3K56me2
H3K56me3
H3K56ac
H3K79me1
H3K79me2
H3K79me3
H3K79ac
H3K122ac

**Figure S2 EpiProfiler confirms IR mediated changes to histone PTMs**

a) Select H3 and H4 residue PTM data from the EpiProfiler Histone PTM dataset are shown. Height of the bars represents the percent of the residue modified at the indicated timepoint. Error bars show standard deviation for three replicates. Significance was determined by a Friedman test comparing timepoints within a given residue. Significance values are as follows: ns $p>0.05$; * $p<0.05$; ** $p<0.01$; *** $p<0.001$; **** $p<0.0001$.

a

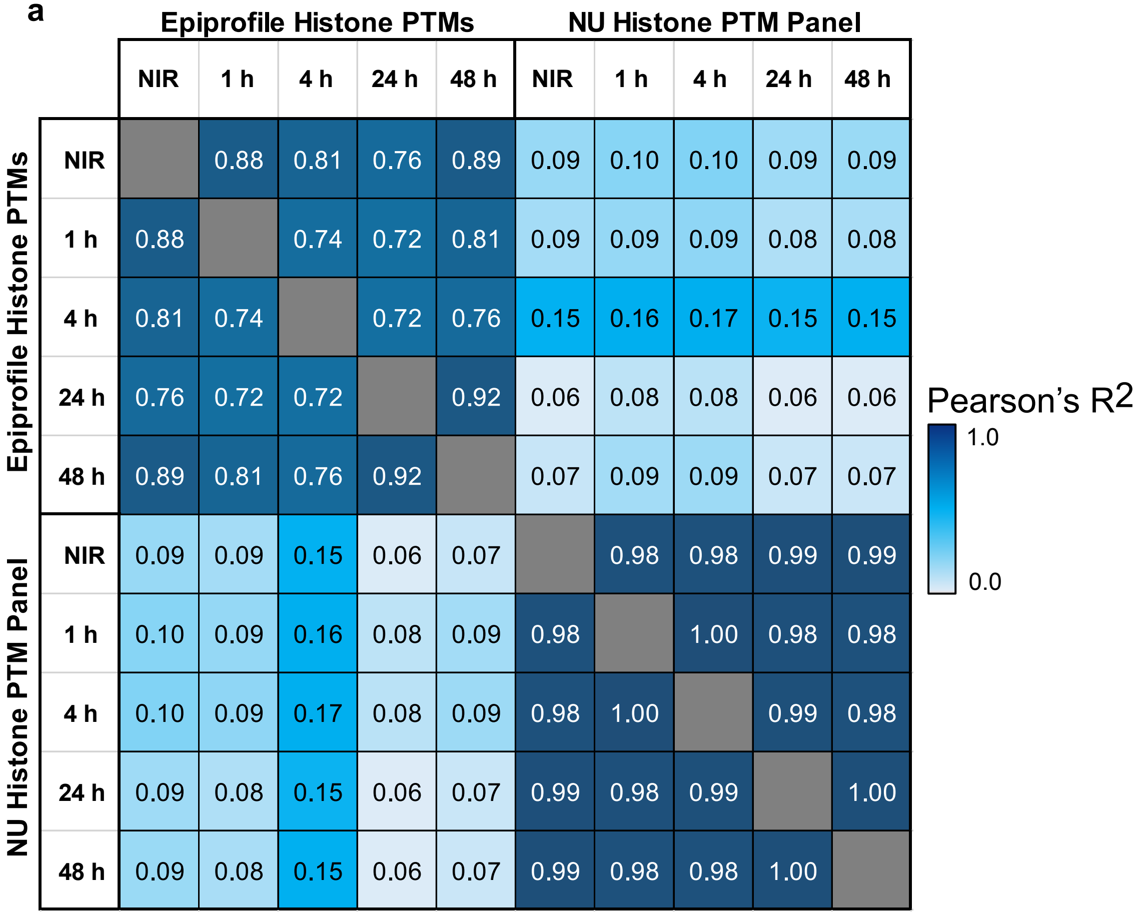
Epiprofile Histone PTMs
NU Histone PTM Panel
NIR 1 h 4 h 24 h 48 h NIR 1 h 4 h 24 h 48 h
Epiprofile Histone PTMs
NIR 0.88 0.81 0.76 0.89 0.09 0.10 0.10 0.09 0.09
1 h 0.88 0.74 0.72 0.81 0.09 0.09 0.09 0.08 0.08
4 h 0.81 0.74 0.72 0.76 0.15 0.16 0.17 0.15 0.15
24 h 0.76 0.72 0.72 0.92 0.06 0.08 0.08 0.06 0.06
48 h 0.89 0.81 0.76 0.92 0.07 0.09 0.09 0.07 0.07
NU Histone PTM Panel
NIR 0.09 0.09 0.15 0.06 0.07 0.98 0.98 0.99 0.99
1 h 0.10 0.09 0.16 0.08 0.09 0.98 1.00 0.98 0.98
4 h 0.10 0.09 0.17 0.08 0.09 0.98 1.00 0.99 0.98
24 h 0.09 0.08 0.15 0.06 0.07 0.99 0.98 0.99 1.00
48 h 0.09 0.08 0.15 0.06 0.07 0.99 0.98 0.98 1.00
Pearson’s R2
1.0
0.0

**Figure S3 EHMP and Epiprofile analyses report differential histone alterations**

a) Matrix shows the correlation between 55 PTMs measured by both EpiProfile and the EHMP assay. Color of the squares is proportional to Pearsons correlation coefficient. The $R^2$ value between two timepoints is shown within each square. Data used to construct the matrix are the average percent residue modification values.

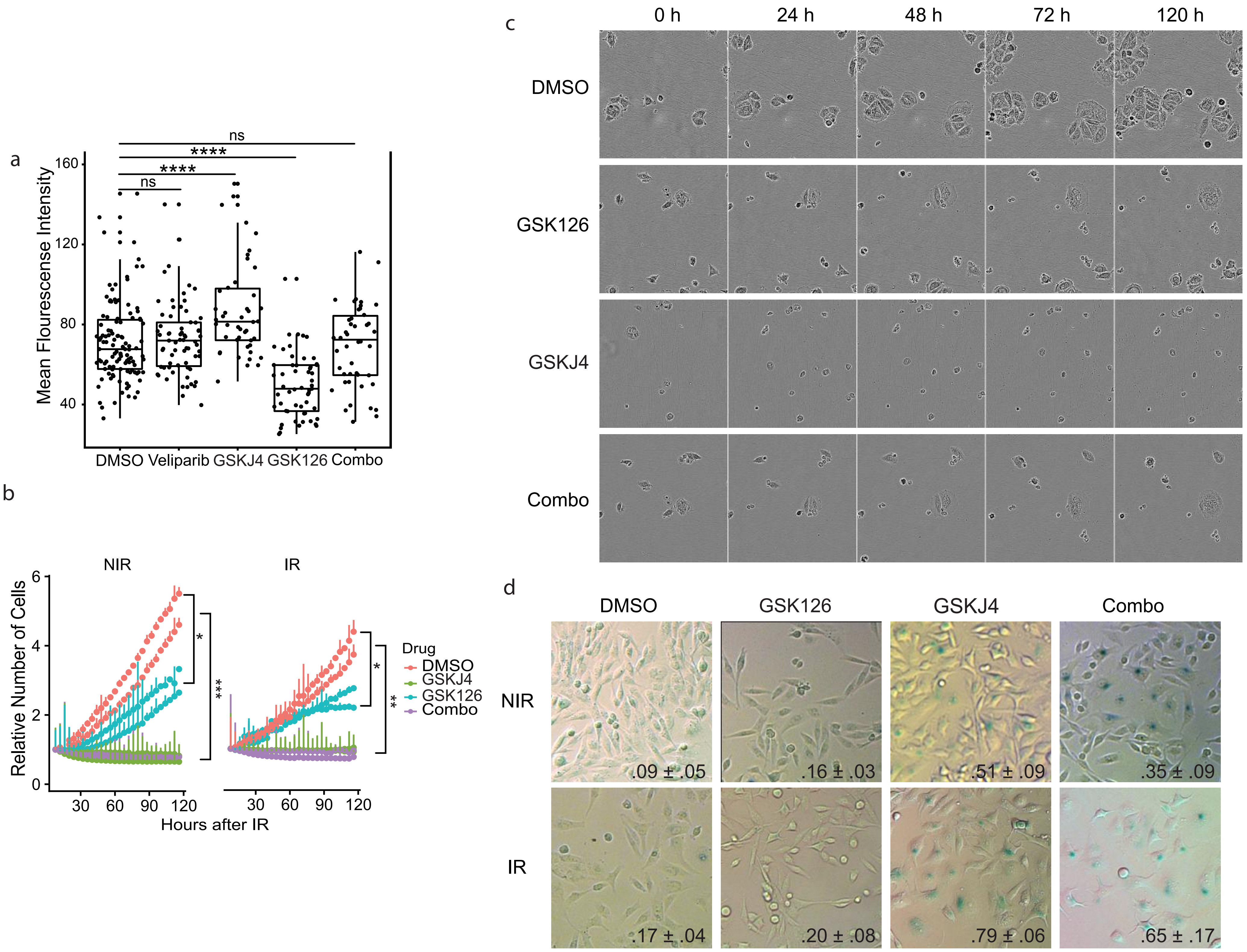

a
Mean Flourescense Intensity
160
120
80
40
ns
****
****
ns
DMSO Veliparib GSKJ4 GSK126 Combo
b
NIR
IR
Relative Number of Cells
6
4
2
0
30 60 90 120
Hours after IR
*
***
**
Drug
DMSO
GSKJ4
GSK126
Combo
c
0 h
24 h
48 h
72 h
120 h
DMSO
GSK126
GSKJ4
Combo
d
DMSO
GSK126
GSKJ4
Combo
NIR
IR
.09 ± .05
.16 ± .03
.51 ± .09
.35 ± .09
.17 ± .04
.20 ± .08
.79 ± .06
.65 ± .17

**Figure S4 Inhibition of H3K27 methylation sensitizes cells to Ionizing Radiation**

a) H3K27me3 mean fluorescent intensity of drug treated cells. Cells were treated and imaged as in Fig 3b. MFI is calculated for each nucleus using a custom ImageJ macro. Significance was determined by a Wilcox Ranked-Sum Test against DMSO treatment. Significance values are as follows: ns $p>0.05$; * $p<0.05$; ** $p<0.01$; *** $p<0.001$; **** $p<0.0001$. Three biological replicates were collected. Total number of datapoints are as follows: 110, 62, 43, 49, 38 per group from left to right.

b) Incucyte growth curves of drug treated cells. Cells were treated for 60 min as in B and then exposed to IR or mock irradiated (NIR). Cell number was tracked for 120 h in an Incucyte system. The mean normalized number of cells is plotted, and error bars denote SEM for 3 replicates. Significance was determined by Dunnett's Multiple Comparisons Test against DMSO treatment.

c) Images excerpted from the Incucyte image dataset over the course of the 120 h analysis. Timepoints are equivalent to Fig 3d. Only the IR condition is shown.

d) SA-βGal staining of cells treated for 1 h with the indicated drugs prior to IR insult and allowed to recover for 72 h before fixation and staining. A representative image, selected from three replicates, is shown for each treatment.
